# Brain and kidney Extracellular Vesicle Morphology and Complementary Brain proteomics after Inflammatory Challenge

**DOI:** 10.64898/2026.10.29.762988

**Authors:** Kent Brown, Yingjie Wang, Zhiying Shan, Lanrong Bi

**Author notes:** Equal contribution (Kent Brown, Yingjie Wang).

## Abstract

Shape, size variation, and protein composition describe different properties of tissue extracellular vesicle (EV) preparations. We compared brain and kidney EV shape and size using transmission electron microscopy (TEM). We also assessed repeated protein detection and reanalyzed a deposited brain proteomics cohort, with attribution to its original source. The study used intracerebroventricular lipopolysaccharide (LPS) and an intravenous miRNA-associated EV formulation. Each group contained three biological samples, as labelled in the source data. Source morphometry summaries included 385 brain and 101 kidney profiles. Kidney circularity was 0.9225, 0.8977, and 0.9288 in the control, LPS, and treatment groups, respectively. The corresponding values for diameter quartile dispersion, a measure of size variation, were 0.2603, 0.5334, and 0.6018. Thus, near-control outline shape coexisted with persistent size variation. None of the six primary morphology tests remained significant after Holm correction. Independent recalculation confirmed 1,527 eligible rat protein accessions across individual searches. The complete quantitative matrix contained 953 features. The main peptide filter combined unique and razor peptides. A stricter analysis retained 916 features. PTGFRN and LAMB2 remained the only significant treatment-versus-LPS findings. In the main analysis, PTGFRN increased 14.71-fold (q = 0.0365), and LAMB2 increased 6.23-fold (q = 0.00585). Their LPS levels were already close to control values. Exploratory normalization used the geometric mean of four measured EV-associated markers. The resulting treatment-versus-LPS increases were 16.43-fold and 6.96-fold, respectively. These ratios support selective changes in relative protein representation within the recovered mixture. They do not establish protein copies per EV. Analysis of the full protein profile and calibration of the complete selection process did not support an overall return to control. This study compares morphology and composition as separate outcomes. It identifies membrane and matrix associations as priorities for validation. Original TEM measurements were unavailable, sample matching remained unresolved, and kidney proteomics were absent. These limitations prevent causal conclusions and individual-sample comparisons between morphology and protein composition.

**Significance Statement:** This study contributes to exosome research by integrating particle morphology and proteomic analysis to characterize tissue-derived extracellular vesicle-enriched preparations following inflammation and miRNA-associated vesicle administration. The findings highlight that particle shape, size heterogeneity, and protein composition can follow different treatment-associated patterns, supporting their assessment as complementary features of exosome preparations. Selective enrichment of PTGFRN and LAMB2 prioritizes membrane and extracellular-matrix association for further investigation. This approach strengthens the interpretation of exosome characterization data and defines focused questions about particle identity, protein localization, and functional cargo delivery. It provides a foundation for evaluating how changes in recovered vesicle preparations relate to the biological activity of exosome-based interventions.

## 1. Introduction

Extracellular vesicle (EV) preparations recovered from tissues provide a molecular and morphological view of material released by, associated with, or co-separated from the source tissue. Interpreting that view requires attention to what each measurement captures. Proteomics estimates relative protein abundance in the recovered mixture. Negative-stain transmission electron microscopy (TEM) records two-dimensional particle profiles after adsorption, staining and drying. Identification lists describe which protein accessions received sufficient evidence in a particular analysis. These observations are complementary, but none alone establishes vesicle integrity, cellular origin or biological activity [1,3].

Our deposited bioRxiv report characterized brain EV-enriched proteomics after intracerebroventricular LPS and subsequent intravenous administration of a miRNA-associated EV formulation [2]. It identified selective protein responses without demonstrated overall convergence toward control. The present manuscript extends that work with brain and kidney morphometry, identification-recurrence analysis and additional quantitative sensitivities. The combined LFQ workbook and nine individual identification files were recalculated for this revision. The brain cohort and its principal differential-abundance results are reused and are not an independent validation cohort. The added scientific question is whether a preparation can show near-control projected outlines while retaining distributional and compositional differences.

Two distinctions guide this extension. First, average outline shape and the width of a diameter distribution answer different questions: near-control circularity can coexist with persistent size dispersion. Second, an accession missing from an individual DDA identification list may still have a measurable quantitative ion signal. DDA incompletely samples complex peptide mixtures, and identification consistency cannot be substituted for differential abundance [22,23]. Separate analysis of particle distributions and identification evidence can therefore reveal limitations that group-average morphology or protein fold changes alone would miss.

A further issue concerns apparent molecular recovery. The same LPS mean enters the LPS/control and treatment/LPS contrasts with opposite signs, allowing sampling error and selection on the first contrast to favor opposing trajectories even without a treatment effect. Relative abundance references also influence effect estimates and uncertainty. We therefore compared animal-level morphology summaries, identification recurrence, protein-specific trajectories and full-profile geometry. Additional peptide-evidence and protein-ratio analyses ask whether the leading compositional signals depend on shared peptides or a common sample-wide intensity scale. This design separates a candidate biological interpretation from the measurement assumptions required to support it.

## 2 Materials and methods

### 2.1 Animals

Adult male Sprague–Dawley (SD) rats were purchased from Charles River Laboratories (Wilmington, MA, USA) and acclimated to the animal care facility for at least one week before experiments. Rats were housed at 20–24 °C under a 12 h light–dark cycle with ad libitum access to food and water. Rats received intracerebroventricular (ICV) injections of LPS or vehicle, with or without intravenous (IV) administration of miRNA, as described below. Brains and kidneys were subsequently collected for EV isolation. All experimental protocols were approved by the Institutional Animal Care and Use Committee at Michigan Technological University.

### 2.2 Study design and data provenance

Adult male SD rats aged 10–12 weeks (200–350 g) were used. They were divided into three experimental groups (n=3 per group), and received saline vehicle control (SD), LPS alone (LPS), and LPS plus a miRNA-associated EV formulation (Mi; hereafter, treatment)—each contributed three independent, unpooled brain preparations [2]. LPS from *Escherichia coli* O26 was administered as a single ICV injection into a lateral ventricle at a dose of 5 µg in 2 µL; control rats received 2 µL saline. Following recovery, rats in the treatment group received a single 20 µL injection of the formulation via the tail vein. Brains were collected 24 h after LPS or saline administration.

The brain TEM images and proteomics data were derived from the same set of preparations. In the label-free quantification (LFQ) dataset, columns F1–F3 correspond to LPS, F4–F6 to treatment, and F7–F9 to control. These labels establish group membership but do not identify the correspondence between individual TEM and LFQ samples within each group. The kidney EV dataset comprised nine separately identified samples from Sprague–Dawley rats in the same three experimental groups, with three samples per group: control, LPS alone, and LPS plus the miRNA-associated EV formulation.

### 2.3 Formulation and tissue EV enrichment

Untreated-brain EV material was combined with hsa-miR-146a-5p, hsa-miR-10a-5p and hsa-miR-21-5p using Exo-Fect. Each reaction contained 10 pmol of each RNA, 15 µL Exo-Fect, 30 µL miRNA mixture, material equivalent to 300-500 µg EV-preparation protein and PBS to 225 µL. After incubation, 45 µL ExoQuick-TC was added; recovered material was resuspended in 120 µL PBS, of which 20 µL was administered per animal [2]. Reaction inputs and injected volume do not establish retained RNA copies, encapsulation or particle-normalized active dose. Treatment associations refer to the complete formulation.

Brain tissue excluding the cerebellum was processed at approximately 0.2 g per 2 mL Hibernate E medium with collagenase D at 2 mg/mL and DNase I at 40 U/mL for 30 min at 37 °C. Straining, sequential centrifugation at 300, 2,000 and 10,000 × g, and 0.22 µm filtration preceded a 30% sucrose-cushion centrifugation at 100,000 × g for 90 min. The pellet was washed by a second 100,000 × g centrifugation and resuspended in 50 µL DPBS [2]. The kidney enrichment using similar protocol to brain EVs preparation. Kidney TEM comparisons used the recorded control, LPS and treatment group labels.

### 2.4. Imaging characterization and morphometry

Representative negative-stain transmission electron microscopy (TEM) composites were available for brain and kidney EV-enriched preparations. The brain immunoblot panel included GM130, CD9, TSG101, and ALIX, with lanes labelled SD, SD+LPS, and SD+LPS+miRNA (Figure S).

ImageJ measurements included area, perimeter, fitted-ellipse axes, circularity, maximum Feret diameter, and related parameters [7]. All 488 original measurement rows were preserved in the audit record. Two brain rows that duplicated all 20 numeric measurements of earlier records were excluded from analysis, leaving 486 profiles: 385 brain and 101 kidney profiles. Equivalent circular diameter (ECD) was recalculated from traced area, correcting 13 misreferenced formulas in Mi1-F01:

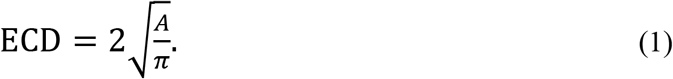

Aspect ratio (AR) was defined as the fitted-ellipse major-axis length divided by the minor-axis length:

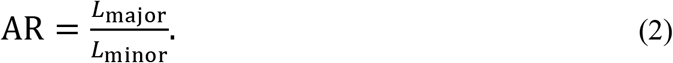

Circularity (*C*) was checked against its geometric definition:

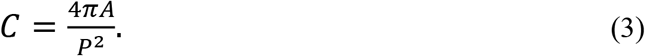

Here, *A* denotes traced area, *P* denotes perimeter, and *L*_major_ and *L*_minor_ denote the fitted-ellipse axis lengths. Small profiles with unverified calibration were retained without a size-based exclusion threshold. Solidity values slightly exceeding one were retained and noted as digitization or export issues.

For the primary analysis, AR and circularity were summarized as the medians of retained profiles pooled across fields within each biological sample. ECD dispersion was calculated separately within each field using the coefficient of quartile dispersion (CQD):

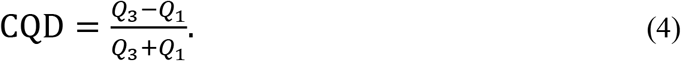

The within-field ECD quartiles, *Q*_1_ and *Q*_3_, were calculated by linear interpolation. Sample-level CQD was the median of the corresponding field-level values. Each group was summarized as the mean ± SD of three equally weighted biological-sample summaries. A sensitivity analysis summarized AR and circularity as the median of the field medians within each sample. Brain samples contributed two to four fields each, whereas each kidney sample contributed one field.

Absolute physical calibration was not verified across fields, particularly for brain LPS1-F02. AR, circularity, and within-field CQD are invariant to uniform isotropic scaling within a field. This property does not eliminate effects of resolution, staining, sample preparation, segmentation, or profile selection. ECD and Feret measurements were retained in their recorded units for auditing and were excluded from physical-size inference.

### 2.5 Identification recurrence and sample accumulation

The nine individual brain protein-identification exports were analyzed separately from the combined label-free quantification (LFQ) matrix. Each export contained 15 fields, including accession, sequence coverage, peptide counts, peptide-spectrum matches (PSMs), unique-peptide counts, protein length, and broad annotations, but no LFQ abundance columns. Records were matched across exports by exact accession, without aggregation by gene symbol.

Only entries explicitly annotated as rat were retained. An accession was excluded from all samples if any export flagged it as a contaminant. This rule excluded A0A8I6AKB2/Pfn1, which had inconsistent contaminant flags across exports. Bacterial Lys-C (Q02SZ7) was excluded by species annotation. The original flags were preserved in the audit record.

Let *S_s_* denote the retained accession set for preparation *s* . For accession *i*, the binary identification indicator was defined as

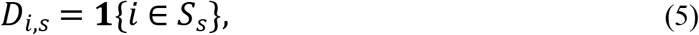

where the indicator equals 1 when accession *i* was identified after filtering in preparation *s* and 0 otherwise. For group *g*, containing three preparations, the identification count was

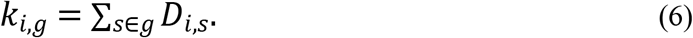

A zero indicator denotes the absence of a retained identification in an export; it does not establish biological absence or zero quantitative abundance.

The group union was defined as *U_gg_* = ⋃*_s_*_∈*gg*_*S_s_*. The fraction of this union identified in all three preparations was calculated as

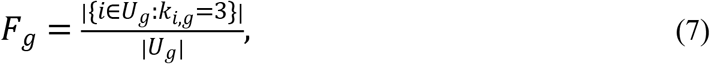

where vertical bars denote the number of elements in a set. Pairwise exact-accession Jaccard similarity was calculated as

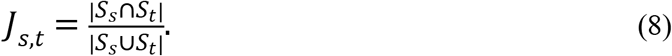

The three within-group pairwise similarities were summarized descriptively because they share preparations. A sensitivity analysis required at least three unique peptides for an identification within each run. This per-run criterion was distinguished from the combined-workbook *Full Filter*, which uses unique plus razor peptides.

The mean accumulated number of identified accessions across all subsets of *m* preparations was calculated exactly as

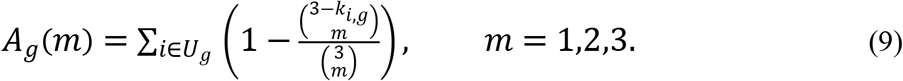

Here, 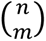 denotes a binomial coefficient and equals zero when *n* < *m*. This calculation averages the identification union over every subset of the specified size. It describes sample accumulation within the observed cohort, incorporating both analytical and biological variation, and does not extrapolate to instrument saturation or the complete tissue proteome. Accession grouping in the individual searches was not assumed to be identical to grouping in the combined LFQ analysis.

### 2.6 Quantitative proteomics and relative references

For each EV-enriched sample, a 10 µg protein aliquot was buffer-exchanged into 25 mM ammonium bicarbonate and digested with trypsin and Lys-C. A single 5 µL injection per sample was analyzed by LC-MS/MS using a two-hour data-dependent acquisition on a Q Exactive HF-X, followed by a blank. Database searching used Proteome Discoverer 3.3.0 with SEQUEST and the rat reference proteome UP000002494, supplemented with enzyme and contaminant sequences. Percolator was used to assess peptide-spectrum matches.

Abundance normalization used the facility’s “Specific Protein Amount” setting. The resulting normalized abundance columns were used for quantitative analysis; vendor ratio columns and the nine individual identification lists were not used as quantitative inputs. No additional missing-value imputation was performed. The facility report did not document upstream missing-value handling or peptide-matching settings.

The combined export contained 1,404 protein groups, of which 978 passed the workbook’s cached *Full Filter*. Inspection of the underlying formulas showed that this filter excluded flagged contaminants, keratins, and immunoglobulins and required a combined count of at least three unique plus razor peptides. This criterion differed from the facility report’s description of at least three unique peptides. Excluding bacterial Lys-C (Q02SZ7) left 977 rat entries. Of these, 953 had positive normalized abundance in all nine samples and formed the primary complete-case matrix. The remaining 24 incomplete entries were excluded without additional imputation.

Among the 953 complete entries, 37 had fewer than three unique peptides. Restricting the matrix to entries with at least three unique peptides therefore yielded a stricter 916-feature sensitivity set. Eligibility for the primary analysis was defined by the workbook formula. Exact accession identifiers were retained without aggregation of entries sharing a gene symbol.

Normalized abundances were log2-transformed:

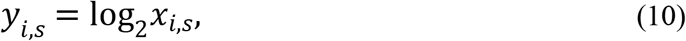

where *x_i_*_,*s*_ is the normalized abundance of feature *i* in sample *s*. Two-sided Welch tests were performed for LPS/control, treatment/LPS, and treatment/control. Benjamini-Hochberg (BH) adjustment was applied separately across the 953 features within each contrast [5]; the three contrast-specific testing families were not pooled. The log2 fold change for groups *g* and ℎ was calculated as

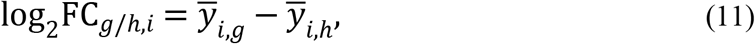

where the overbars denote group means. Fold changes therefore represent ratios of group geometric means. Nominal candidates required *p* < 0.05 and |log_2_FC| ≥ log_2_(1.5) . This nominal screen did not control the false discovery rate. The same contrasts were recalculated in the stricter 916-feature set, with BH adjustment performed separately within each contrast across those 916 features.

Principal component analysis (PCA) was performed after centering each protein across samples, without variance scaling. Exploratory global multivariate tests enumerated all 1,680 assignments of the nine whole sample columns to three labelled groups of three. These analyses reproduce and extend the original cohort analysis [2].

Facility-normalized abundances remained the primary quantitative reference. A median-centering sensitivity analysis subtracted each sample’s median log2 abundance across the fixed set of 953 features and added a common grand-median value [8]:

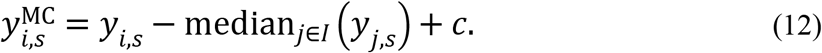

A fixed-feature centered-log-ratio (CLR) sensitivity analysis subtracted each sample’s mean log2 abundance across exactly the same 953 features [19]:

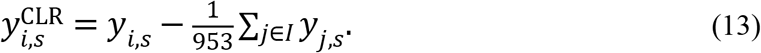

Here, *I* is the fixed 953-feature set and *c* is the common grand-median value. All input abundances were positive, so no pseudocount was added. CLR provided an alternative relative reference without assuming that LFQ intensities constituted an exactly measured closed mass composition. Contrasts, BH adjustment, and centroid geometry were evaluated under each reference. Additional effect-estimation details are provided in Supplementary Methods S1.

An additional descriptive ratio analysis used four measured EV-associated proteins as alternative references: CD9 (A0A8I5ZZU3), ALIX/PDCD6IP (Q9QZA2), FLOT1 (Q9Z1E1), and FLOT2 (A0A0G2K0Z1). Within each sample, log2 ratios for PTGFRN and LAMB2 were calculated against each marker separately and against the arithmetic mean of the four marker log2 intensities. For the four-marker panel, this was calculated as

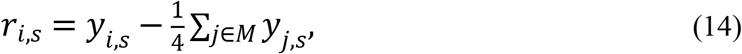

where *M* denotes the four-marker reference panel. This calculation is equivalent to using the geometric mean of the four marker intensities as the reference on the original abundance scale. Treatment/LPS fold changes were calculated from the difference between group mean log2 ratios:

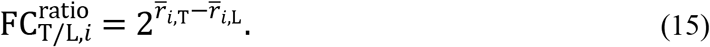

Here, T denotes treatment and L denotes LPS. Any common multiplicative factor applied to all intensities within a sample cancels exactly in these ratios. The marker panel was selected during secondary analysis, and marker abundance per EV was not assumed to be stable. Ratios were reported descriptively, without an additional significance screen. They describe relative protein representation in the recovered material and do not establish molar stoichiometry or protein copies per particle.

### 2.7 Shared reference calibration and statistical inference

For protein *i*, let *y_i_*_,*gg*_ denote the mean log2 abundance in group *g*, with C, L, and T denoting control, LPS, and treatment, respectively. The LPS/control and treatment/LPS contrasts were defined as

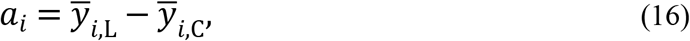

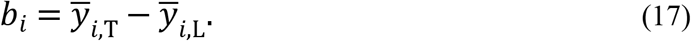

The treatment/control contrast was therefore

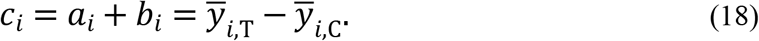

Opposing movement was defined by *a_i_b_i_* < 0, whereas closer proximity to control required |*c_i_*| < |*a_i_*|. Because both contrasts share the LPS group, their sampling errors are negatively correlated when group sampling errors are independent:

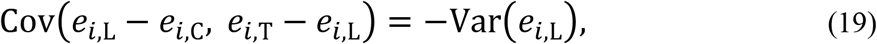

where *e_i_*_,*gg*_ is the sampling error of the corresponding group mean. This identity concerns sampling errors for an individual protein, not biological covariance across proteins. Candidate selection based on *a_i_* can further accentuate apparent reversal arising from the shared LPS reference.

Conditional calibration used a reference of no difference between the LPS and treatment groups. The three original control columns were held fixed, and the six LPS/treatment columns were reassigned through all 20 allocations into two groups of three. Entire sample columns were reassigned together, preserving the observed dependence among proteins. For each allocation, the LPS/control Welch screen was repeated using the same nominal and fold-change criteria (*p* < 0.05 and |log_2_FC| ≥ log_2_(1.5)). The fractions showing opposing movement and closer proximity to control were recalculated within the newly selected candidate set. Whole-profile geometry was also recomputed using the fixed set of 953 features.

Let *z_π_* denote a diagnostic statistic under allocation *π*, *z*_obs_ its observed value, and *Π* the set of 20 allocations, including the observed allocation. Inclusive upper- and lower-tail reference fractions were calculated as

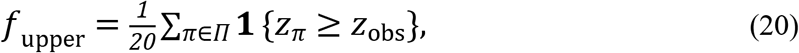

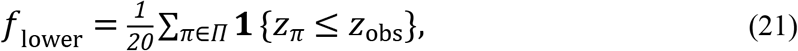

where the indicator equals 1 when the stated condition is met and 0 otherwise. These fractions have a one-sided resolution of 0.05. The calibration diagnostics were not treated as a confirmatory family of multiple statistical tests. Their interpretation requires exchangeability of the six LPS/treatment columns; equality of group means alone is insufficient when the underlying distributions differ.

Whole-profile geometry used contrast vectors **a** and **b**, each containing the corresponding contrasts for all 953 features. The reversing projection coefficient was calculated as

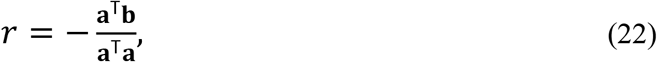

where the superscript T denotes transpose. The component orthogonal to the LPS/control contrast was

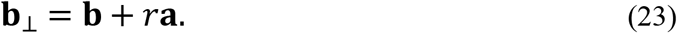

Its fraction of the squared treatment/LPS contrast length was

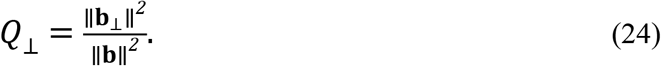

The relative reduction in Euclidean distance to control was calculated as

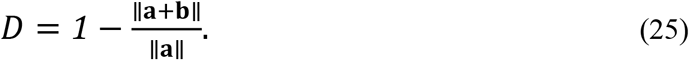

Positive *D* indicates a shorter distance to control, whereas negative *D* indicates a greater distance. These quantities describe contrast geometry and do not represent proportions of proteins rescued.

Bootstrap uncertainty was evaluated across all 19,683 ordered resampling configurations, drawing three whole sample columns with replacement within each group. Percentile intervals were exploratory given three biological samples per group. Column omissions and all 27 balanced two-per-group subsets assessed within-cohort influence without adding biological replication.

Omnibus TEM tests used the ANOVA F statistic over all 1,680 allocations of nine biological-sample summaries to three labelled groups of three. Holm adjustment covered six tests: AR, circularity, and CQD in each tissue [6]. Planned LPS/control and treatment/LPS contrasts each enumerated all 20 allocations of six sample summaries to two groups of three. Holm adjustment covered 12 planned comparisons: two tissues, three endpoints, and two contrasts.

For a symmetric, nonrandomized, two-sided three-versus-three allocation test, complementary assignments give a minimum attainable *p* value of 0.10. TEM tests required group-label exchangeability; equal means alone were insufficient under unequal distributions. Profiles and fields were not treated as independent animals.

Exploratory log-quartile asymmetry and specimen-assignment analyses are described in Supplementary Methods S1. Individual protein-morphology associations were not inferred without a verified sample crosswalk. Functional-module balances and protein-ratio validation were not performed and remain future work.

## 3 Results

### 3.1 Source characterization and measurement coverage

Negative-stain TEM showed heterogeneous particle profiles in brain and kidney preparations from all three groups (Figures 1 and 2). These images characterize material adsorbed to the grid and do not establish particle yield, purity, or biological activity. The brain immunoblot panel is presented separately as source characterization (Supplementary Figure S3). The individual protein-identification exports identified CD9 and ALIX/PDCD6IP in seven of nine preparations each. Calnexin/CANX was identified in all nine preparations, with 14-23 unique peptides per preparation. Thus, EV-associated proteins were recovered alongside proteins associated with intracellular membranes. These findings support the operational designation EV-enriched preparation without establishing a homogeneous vesicle population.

**Figure 1.**
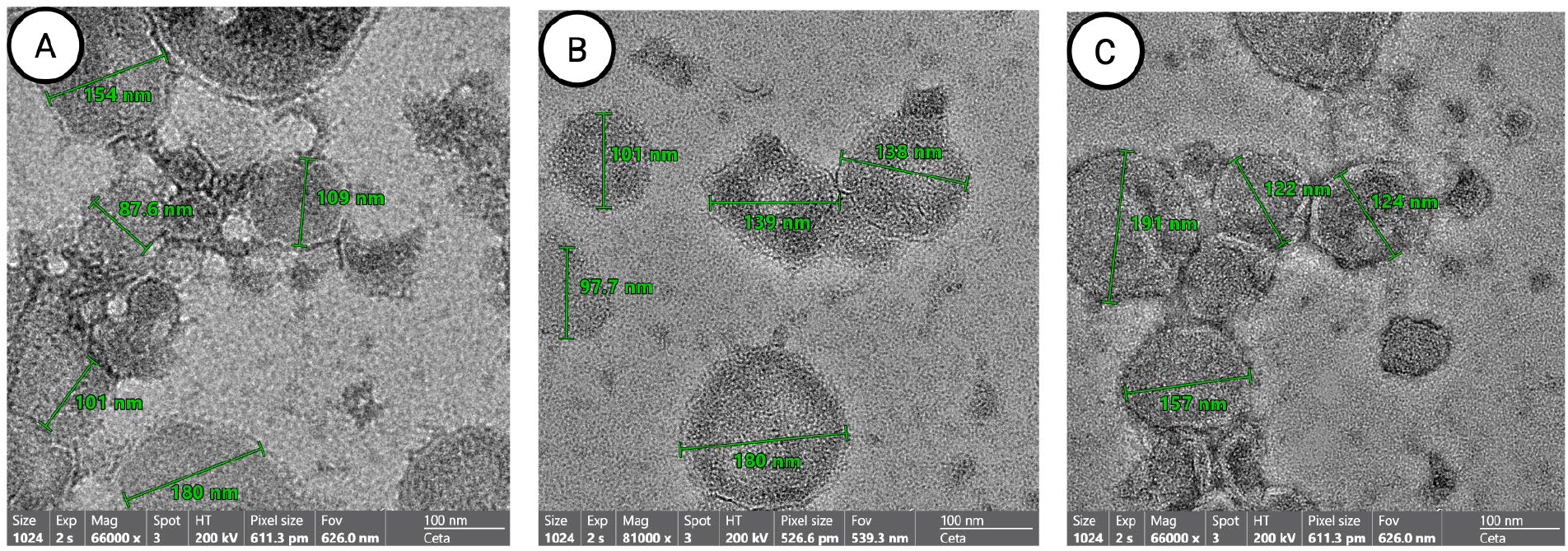
Kidney particle profiles. (A-C) Source negative-stain TEM fields labelled kidney control, LPS and treatment, respectively. Displayed scale bars are 100 nm. Green source-image line annotations are not the area-derived ECD endpoint and are not used for group diameter inference. The marker panel originally duplicated in both tissue composites is shown once in Supplementary Figure S3; its tissue attribution is unresolved.

**Figure 2.**
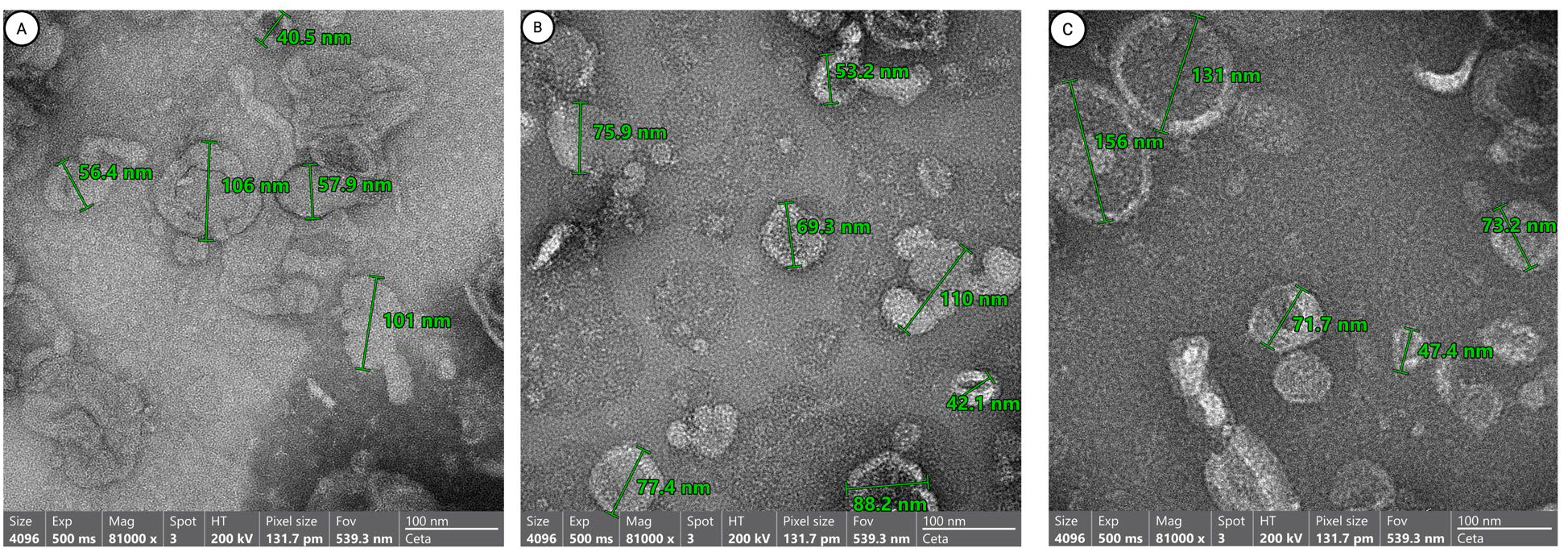
Brain particle profiles. (A-C) Source negative-stain TEM fields labelled brain control, LPS and treatment, respectively. Displayed scale bars are 100 nm. These images provide descriptive context for the source morphometry summaries and do not establish vesicle integrity, particle yield or treatment efficacy. The source-supplied marker panel is presented separately in Supplementary Figure S3.

The source morphometry audit reported 385 retained brain profiles from 27 fields and 101 kidney profiles from nine fields. Control, LPS, and treatment preparations contributed 103, 138, and 144 brain profiles and 39, 28, and 34 kidney profiles, respectively. The source audit excluded two duplicate brain rows and corrected 13 ECD formulas. Each tissue retained three biological preparations per group. The original profile tables were unavailable for independent recalculation; the morphometric values and tests reported below therefore reflect the source analysis. Profile counts describe sampling within the selected fields and cannot estimate tissue EV production or concentration.

Absolute diameter comparisons were not supported across the exported fields. For example, brain LPS1-F02 had a median ECD of 378.94 in recorded units, compared with 51.58 and 48.23 in the other fields from that sample. Resolving this discrepancy requires the original image-calibration and processing records. The earlier report’s nanometer-scale brain summary is therefore not repeated here [2]. AR, circularity, and within-field CQD are invariant to uniform isotropic scaling within a field, although they remain sensitive to resolution, tracing, and profile selection.

### 3.2 Kidney outline shape and diameter dispersion showed different numerical patterns

Kidney group mean circularity, calculated from three sample-level medians, was 0.9225 ± 0.0174 in control, 0.8977 ± 0.0153 after LPS, and 0.9288 ± 0.0145 after treatment (mean ± SD; Table 1; Figure 3). The treatment/LPS difference was +0.0311, and treatment exceeded control by 0.0063. Mean AR was 1.2607 ± 0.1165, 1.1950 ± 0.0465, and 1.2326 ± 0.0538 in control, LPS, and treatment, respectively. For both outline metrics, the treatment mean was numerically closer to control than the LPS mean was. However, the source allocation tests did not establish a treatment effect after multiplicity adjustment. Circularity is bounded and sensitive to perimeter tracing; these small absolute differences cannot be interpreted as a percentage of structural recovery.

**Figure 3.**
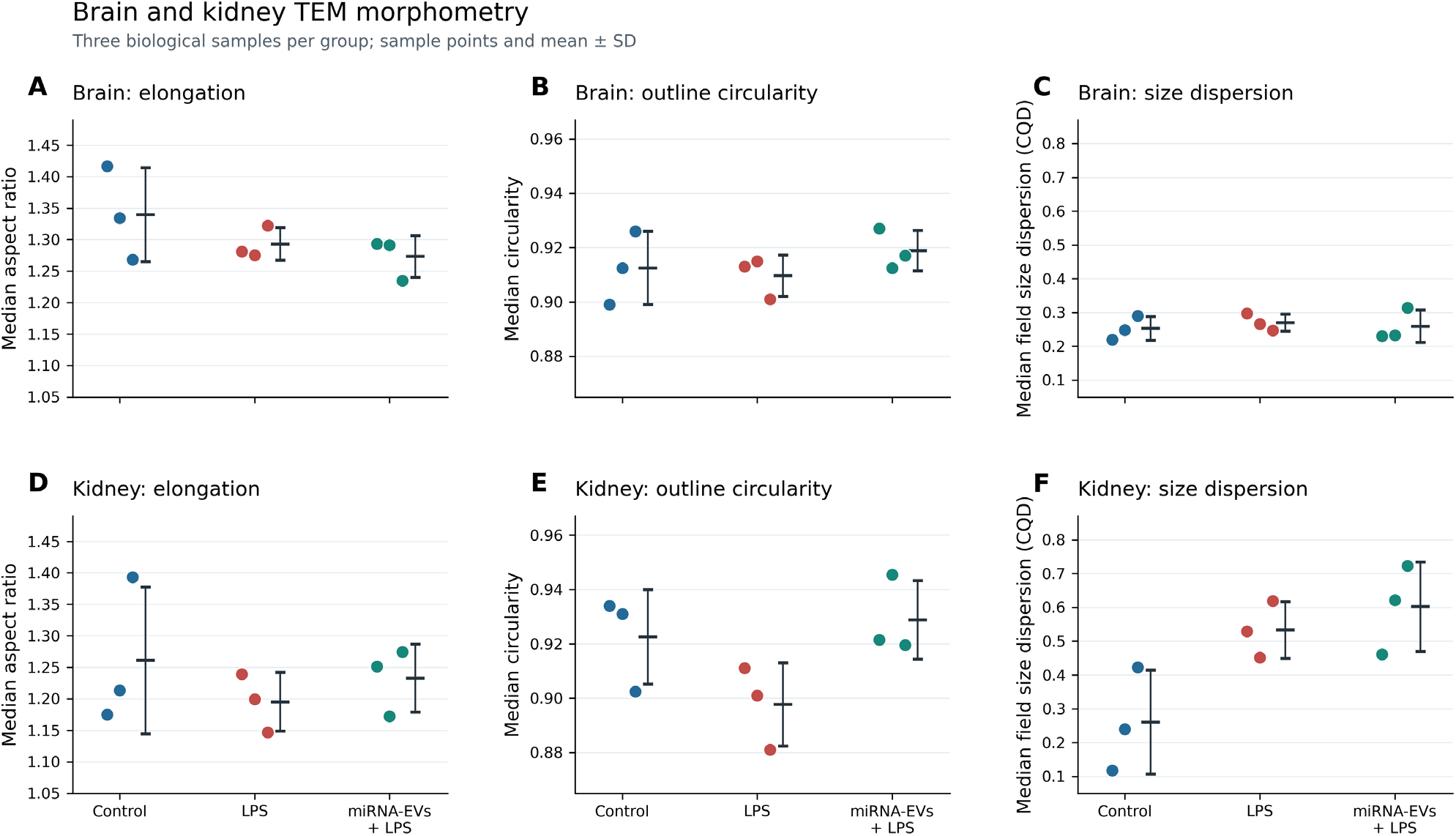
Brain and kidney morphometry at the biological sample level. (A-C) Brain AR, circularity and CQD, respectively. (D-F) Corresponding kidney endpoints across control, LPS and treatment. Each point is one biological sample, n = 3 per group; bars summarize group mean ± sample SD. AR and circularity are medians of retained profiles pooled within a sample. CQD is calculated from area-derived ECD quartiles within each field, then summarized by the median across fields within each sample. Brain samples contribute two to four fields and kidney samples one. The source analysis retained 385 brain and 101 kidney profiles; these are subsamples, not independent biological replicates. None of six primary tissue-endpoint omnibus allocation tests met Holm-adjusted P < 0.05. Numerical changes in circularity and dispersion describe different properties of the sampled profiles.

**Table 1.** Biological sample TEM summaries Mean ± between-sample SD across three source-labelled biological samples per group, retained from the source morphometry analysis. The original row-level export and ROIs were unavailable for independent recalculation in this revision. AR and circularity are sample medians pooled across retained profiles. CQD is the median of field-specific (Q3−Q1)/(Q3+Q1) values of area-derived ECD. Brain samples contribute two to four fields and kidney samples one. All endpoints are dimensionless; profile counts do not enlarge biological n.

| Tissue and group | AR | Circularity | Field CQD |
| --- | --- | --- | --- |
| Brain Control | 1.3396 ± 0.0744 | 0.9125 ± 0.0135 | 0.2527 ± 0.0354 |
| Brain LPS | 1.2928 ± 0.0257 | 0.9097 ± 0.0076 | 0.2701 ± 0.0253 |
| Brain Treatment | 1.2731 ± 0.0332 | 0.9188 ± 0.0074 | 0.2592 ± 0.0478 |
| Kidney Control | 1.2607 ± 0.1165 | 0.9225 ± 0.0174 | 0.2603 ± 0.1537 |
| Kidney LPS | 1.1950 ± 0.0465 | 0.8977 ± 0.0153 | 0.5334 ± 0.0837 |
| Kidney Treatment | 1.2326 ± 0.0538 | 0.9288 ± 0.0145 | 0.6018 ± 0.1322 |

Diameter dispersion showed a different pattern. Kidney CQD increased from 0.2603 ± 0.1537 in control to 0.5334 ± 0.0837 after LPS and 0.6018 ± 0.1322 after treatment. The LPS/control and treatment/LPS differences were +0.2731 and +0.0684, respectively, leaving treatment +0.3415 above control. The treatment group mean CQD was approximately 2.31 times the control mean. This is a ratio of group summaries, not an animal-level fold change. Mean within-field Q3/Q1 ratios were 1.788, 3.382, and 4.404 in control, LPS, and treatment, respectively, indicating a broader multiplicative span of the central half of the sampled diameter distribution. These ratios re-express the same quartiles and provide no independent test. Proximity to control in the outline summaries therefore coexisted with elevated diameter dispersion; the data did not show a uniform return to the control morphometric pattern.

### 3.3 Brain morphometric differences were small and sensitive to field aggregation

Brain mean sample AR was 1.3396 ± 0.0744, 1.2928 ± 0.0257, and 1.2731 ± 0.0332 in control, LPS, and treatment, respectively. Corresponding circularity values were 0.9125 ± 0.0135, 0.9097 ± 0.0076, and 0.9188 ± 0.0074, and CQD values were 0.2527 ± 0.0354, 0.2701 ± 0.0253, and 0.2592 ± 0.0478 (Table 1). Brain CQD differed by +0.0174 for LPS/control and -0.0109 for treatment/LPS, leaving treatment +0.0065 above control. Mean within-field Q3/Q1 ratios were 1.680, 1.743, and 1.708. The sampled brain preparations therefore showed a smaller numerical range of central diameter dispersion than the kidney preparations. Differences in field coverage and preparation metadata constrain this comparison to the available preparations and prevent inference of a tissue-specific biological interaction.

The brain AR trajectory changed when sample summaries were calculated as the median of field medians. Under this aggregation rule, group means were 1.3166, 1.3356, and 1.2656 for control, LPS, and treatment, respectively. The control-to-LPS direction was therefore reversed relative to the primary pooled-profile sample medians. Corresponding circularity means were 0.8968, 0.9118, and 0.9188. These changes indicate that field aggregation materially influenced the small brain shape differences.

None of the six primary tissue-endpoint omnibus tests met Holm-adjusted *p* < 0.05. Kidney CQD had a nominal allocation *p* value of 0.0214 and an adjusted value of 0.1286; kidney circularity had corresponding values of 0.1071 and 0.5357. All 12 planned pairwise comparisons had adjusted *p* = 1.00. The specified symmetric two-sided three-versus-three allocation test has a minimum attainable *p* value of 0.10, limiting inferential resolution. These findings do not establish equivalence or exclude biologically relevant effects.

An exploratory quartile-asymmetry analysis added distributional description without evidence of a treatment effect after multiplicity adjustment (Supplementary Figure S2B). Brain group means were 0.2043, -0.0923, and -0.0579 in control, LPS, and treatment, respectively; kidney means were 0.1842, -0.1872, and 0.3573. Holm-adjusted omnibus *p* values were 0.2929 for brain and 0.3500 for kidney. Sparse field coverage and sensitivity to profile omission limited interpretation of these estimates.

### 3.4 Identification recurrence differed amid unequal identification depth

The nine individual protein-identification exports contained 1,539 distinct accession identifiers. Restricting the analysis to rat entries and excluding any accession flagged as a contaminant in any run retained 1,527 accessions, including 267 identified in all nine preparations. Eleven accessions were flagged in at least one run, and the unflagged bacterial digestion-enzyme entry was excluded separately. This accession universe differs from the 1,404 protein groups in the combined analysis and the 953 complete quantified features; these counts are not interchangeable measures of coverage.

Group unions contained 1,125, 1,183, and 1,142 accessions in control, LPS, and treatment, respectively. The corresponding three-preparation cores contained 461, 425, and 550 accessions, yielding core fractions of 40.98%, 35.93%, and 48.16%. Mean within-group Jaccard similarities were 0.5586, 0.5101, and 0.6200 (Table 2; Figure 4). Treatment preparations therefore had greater observed identification recurrence. However, identification depth ranged from 578 to 975 accessions in LPS, compared with 739-848 in control and 812-865 in treatment. Retained protein-entry PSM sums also differed between LPS1 and LPS3: 3,647 versus 6,679. Unequal identification depth limits attribution of the recurrence pattern to a biological treatment effect. These PSM sums were calculated across protein entries and are not counts of distinct spectra.

**Figure 4.**
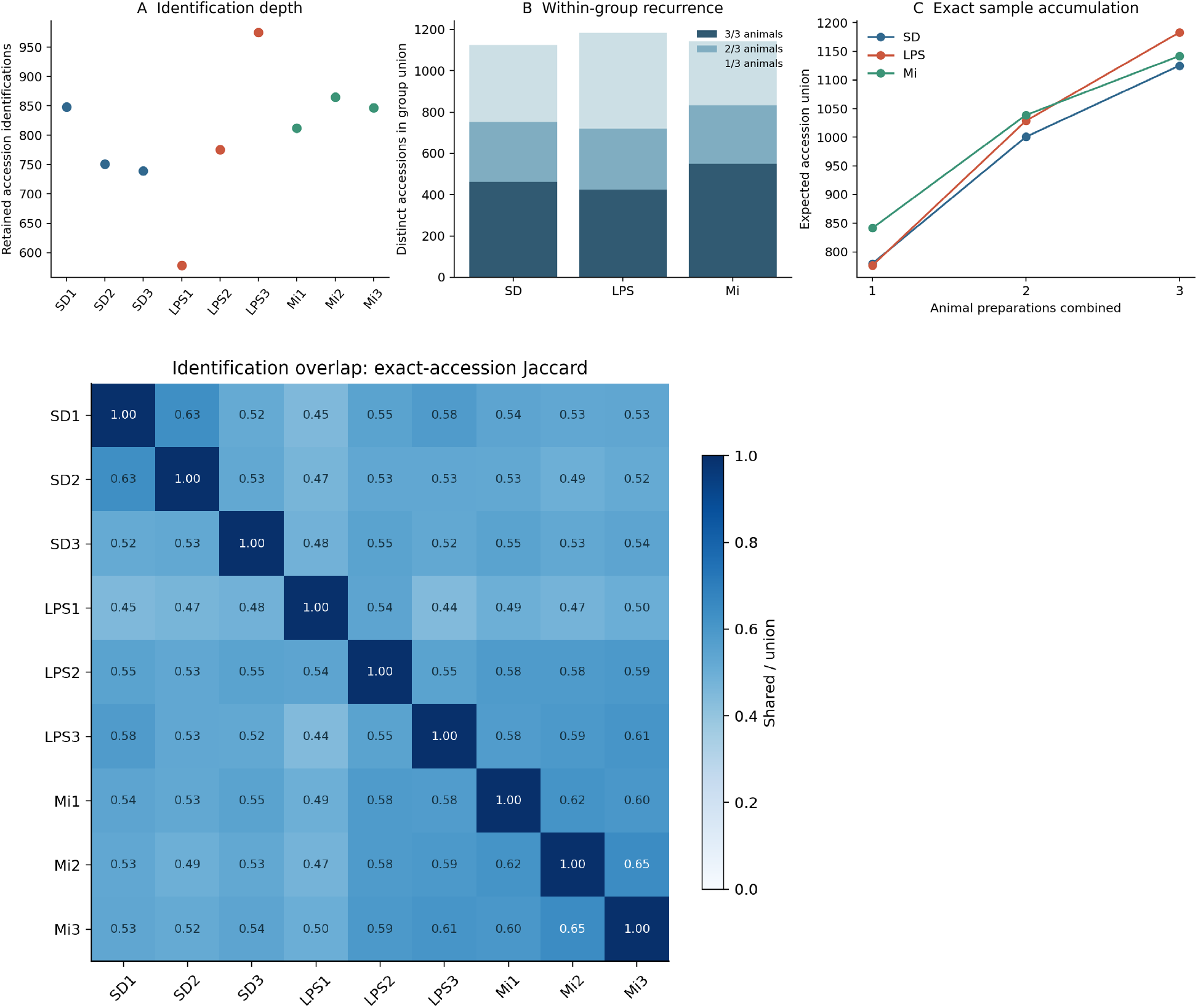
Brain identification depth and recurrence. Upper panels show (A) retained exact-accession identifications per preparation, (B) group unions partitioned by identification in one, two or three preparations, and (C) exact expected identification unions over all subsets of one, two or three preparations. The lower heatmap gives pairwise exact-accession Jaccard similarities. Rat entries were retained after any-run contaminant exclusion, without gene collapsing or quantitative imputation. The three groups contain three preparations each. Overlap values and accumulation points reuse the same samples; they are not independent experiments. Greater recurrence in treatment describes identification consistency, not restored abundance or EV function.

**Table 2.** Brain protein identification recurrence Exact rat accession IDs after conservative any-run contaminant exclusion. Each group has three preparations. Core fraction uses the group union as denominator. Jaccard similarities share preparations and are descriptive. The per-run unique-peptide sensitivity is distinct from the combined LFQ filter.

| Metric | Control | LPS | Treatment |
| --- | --- | --- | --- |
| Union identified in $\geq 1$ of 3 | 1,125 | 1,183 | 1,142 |
| Identified in $\geq 2$ of 3 | 752 | 720 | 832 |
| Core identified in all 3 | 461 | 425 | 550 |
| Core fraction | 40.98% | 35.93% | 48.16% |
| Mean within-group Jaccard | 0.5586 | 0.5101 | 0.6200 |
| Per-preparation ID range | 739-848 | 578-975 | 812-865 |
| Core fraction with $\geq 3$ unique peptides per run | 36.74% | 34.00% | 45.90% |

Requiring at least three unique peptides in each contributing run yielded a 987-accession union and a 161-accession core shared across all nine preparations. Within-group core fractions remained highest in treatment, followed by control and LPS, at 45.90%, 36.74%, and 34.00%, respectively. Exact sample accumulation increased from mean single-preparation counts of 779.33, 776.00, and 841.33 in control, LPS, and treatment to the respective three-preparation unions of 1,125, 1,183, and 1,142. These curves are deterministic summaries of the same preparations and do not add biological replication.

The recurrence audit also distinguished detection evidence from biological absence. CD81 was identified in five of nine individual searches but did not meet the combined three-peptide filter. CD63 and TSG101 were not identified in the individual lists, although TSG101 was included in the labelled immunoblot panel. Such differences do not establish biological absence or invalidate an antibody signal, because data-dependent acquisition and immunoblotting have different detection properties. Conversely, CANX identification in all nine lists prevents using the marker panel alone to conclude that intracellular membrane-associated material was excluded.

The leading quantitative candidates had different identification and quantification coverage. PTGFRN appeared in four individual exports, including controls, whereas the exact LAMB2 accession appeared only in SD3. Both nevertheless had positive normalized LFQ values in all nine columns of the combined workbook. These differences do not, by themselves, establish a processing error: separate searches and the combined analysis may use different protein-inference and signal-extraction procedures. However, the available exports do not establish the peptide-level provenance of every quantitative value. Peptide-level evidence and upstream matching, extraction, and missing-value settings require review before either protein is considered a validated mechanistic marker.

### 3.5 Protein trajectories distinguished relative enrichment from persistent differences

Independent recalculation reproduced the deposited 953-feature analysis [2]. The LPS/control screen contained 227 nominal candidates meeting the 1.5-fold threshold, comprising 95 increases and 132 decreases; none met contrast-specific BH *q* < 0.05. Treatment/LPS contained 56 nominal candidates, comprising 39 increases and 17 decreases, with two BH discoveries.

Treatment/control contained 344 nominal candidates and 174 BH discoveries, comprising 88 increases and 86 decreases (Table 3; Supplementary Figure S1). These counts describe distinct contrast-specific evidence patterns. They do not establish that treatment directly altered 174 proteins, and the absence of LPS/control BH discoveries does not establish an absence of an LPS response.

**Table 3.** Protein contrast counts under three quantitative references All references use the same 953 complete protein groups and separate BH correction within each contrast. The primary nominal screen is P < 0.05 and ≥1.5-fold magnitude. Primary and median-centered results reuse the deposited cohort analysis [2]; CLR is the added fixed-feature sensitivity. References do not create independent data sets.

| Contrast | Primary nominal screen | Primary $q < 0.05$ | Primary $q < 0.05$ and $\geq 1.5$-fold | Median centered $q < 0.05$ | CLR $q < 0.05$ |
| --- | --- | --- | --- | --- | --- |
| LPS/control | 227 | 0 | 0 | 1 | 3 |
| Treatment/LPS | 56 | 2 | 2 | 1 | 0 |
| Treatment/control | 344 | 174 | 163 | 162 | 174 |

PTGFRN (F1M790) and LAMB2 (M0R6K0) were the two treatment/LPS discoveries. PTGFRN increased 14.71-fold (95% pointwise Welch confidence interval, 9.49-22.78; *q* = 0.0365), and LAMB2 increased 6.23-fold (95% pointwise Welch confidence interval, 5.42-7.16; *q* = 0.00585). Their LPS/control ratios were 0.906 and 1.068, and their treatment/control ratios were 13.32 and 6.65, respectively. The leading signals therefore reflected relative enrichment above both comparison groups, rather than reversal of large measured LPS deficits. PTGFRN and LAMB2 had five and seven unique peptides, respectively, in the combined export. Both remained the only treatment/LPS discoveries in the stricter 916-feature analysis, with *q* values of 0.0350 and 0.00562. That analysis yielded 0, 2, and 167 discoveries for LPS/control, treatment/LPS, and treatment/control, respectively. Thus, the two discoveries were retained under the stricter unique-peptide eligibility criterion.

Other proteins illustrated numerical return toward control and persistent differences (Supplementary Table S2). C3 changed from 2.053-fold after LPS to 1.068-fold after treatment relative to control, whereas WFS1 changed from 0.329-fold to 0.919-fold. Their treatment/LPS *q* values were 0.412 and 0.134, respectively. Treatment/control differences remained for MX1 (79.51-fold, *q* = 0.0119), AQP4 (0.137-fold, *q* = 0.0111), SLC1A2 (0.465-fold, *q* = 0.0119), GRIA4 (0.535-fold, *q* = 0.0237), and MFGE8 (0.317-fold, *q* = 0.00840). None of these five proteins had treatment/LPS *q* < 0.05. These selected trajectories distinguish numerical return, residual difference, and treatment-associated relative enrichment; they are not independently tested pathway scores.

A descriptive marker-reference analysis examined whether the leading enrichments tracked a common EV-associated protein reference. PTGFRN treatment/LPS ratios increased 14.80-18.44-fold when referenced separately to CD9, ALIX, FLOT1, and FLOT2; corresponding LAMB2 increases were 6.27-7.82-fold. Using the geometric mean of the four markers as the reference yielded increases of 16.43-fold and 6.96-fold, respectively (Supplementary Table S4). The four-marker reference itself had a treatment/LPS ratio of 0.895. These descriptive ratios are invariant to a common multiplicative sample factor and indicate disproportionate representation of PTGFRN and LAMB2 relative to the measured panel. They do not establish selective sorting, a stable denominator per vesicle, or localization of the proteins on the same individual particles.

Reference choice affected inferential status more than the direction of the large effect estimates. Under median centering, PTGFRN increased 13.01-fold (*q* = 0.0342), whereas LAMB2 increased 5.51-fold (*q* = 0.165). Fixed-feature CLR retained positive estimates of 13.20-fold and 5.59-fold, respectively, but neither protein met *q* < 0.05 (*q* = 0.0697 and 0.178; Figure 5C). CLR discovery counts were 3, 0, and 174 for LPS/control, treatment/LPS, and treatment/control, respectively; its LPS/control discoveries were CANX, SCCPDH, and TMEM30A. These secondary-reference analyses do not replace the primary inference. The treatment/LPS mean-reference shift was 0.156 log2 units under CLR and 0.177 under median centering. Although these shifts were small relative to the leading effects, sample-specific shifts also changed the estimated group variances. Crossing the *q* = 0.05 threshold therefore reflected changes in both relative reference and estimated uncertainty.

**Figure 5.**
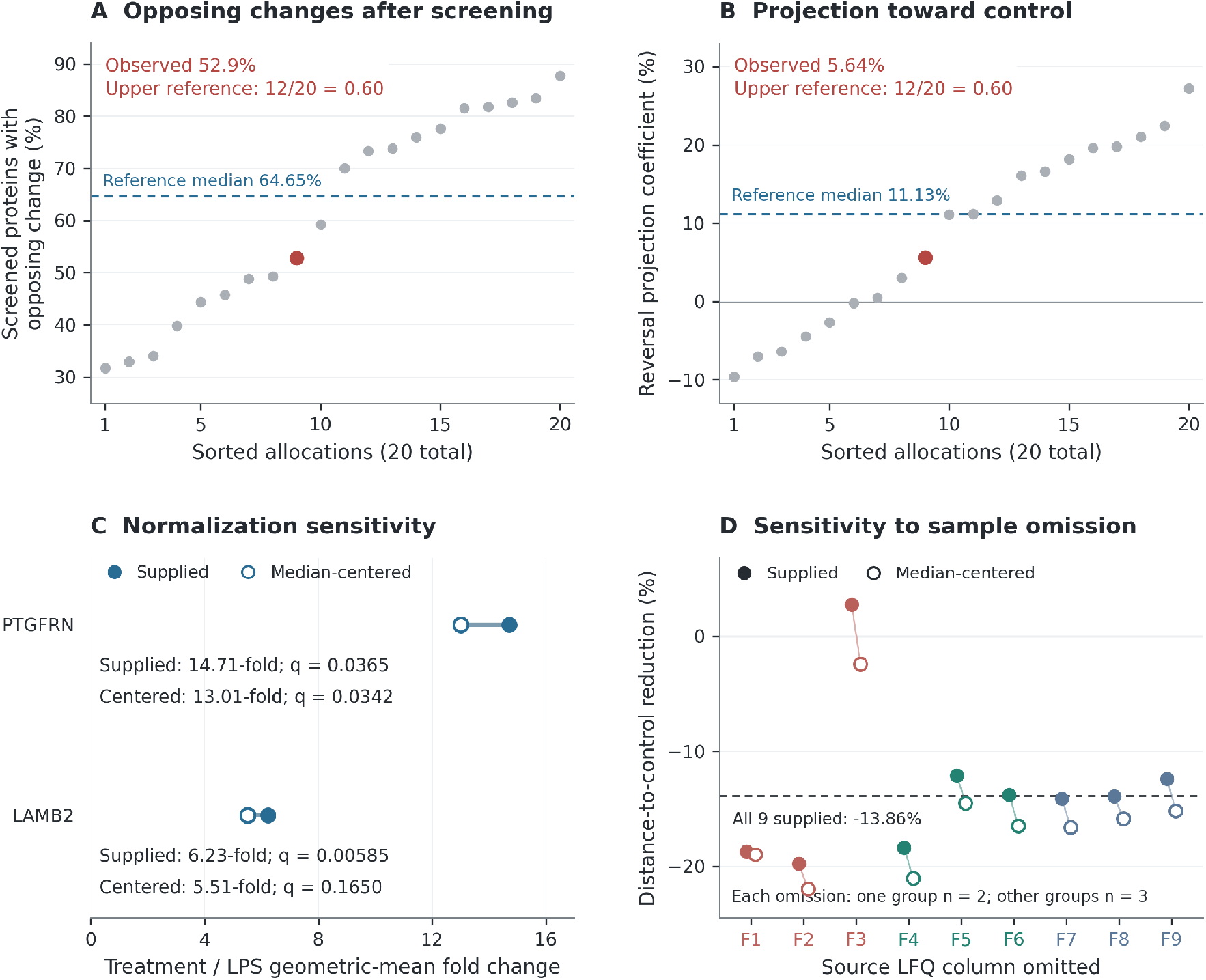
Shared reference calibration and quantitative sensitivity. (A) Opposing fractions after repeating the complete nominal LPS/control candidate screen under all 20 allocations of the six LPS/treatment columns, with controls fixed. Points are allocations, not independent experiments; the observed assignment is highlighted. Its inclusive upper reference fraction is 12/20 = 0.60. (B) Reversing projection coefficient across the fixed 953 proteins under the same allocations. The coefficient describes projection onto the LPS/control vector and is not a percentage of recovered proteins. (C) Previously reported PTGFRN and LAMB2 treatment/LPS fold changes and contrast-specific BH q values under supplied normalization and median centering [2]. Connecting lines imply no uncertainty interval. (D) Previously reported single-source-column omission sensitivities [2]; each omission leaves n = 2 in one group and n = 3 in each other group. The plotted quantity is control-distance reduction under the two references. Omission ranges are not confidence intervals. The added allocation diagnostics require exchangeability and do not measure biological recovery or harm.

Whole-profile differences were evident despite the small number of primary treatment/LPS discoveries. Under the supplied normalization, PC1 and PC2 accounted for 54.51% and 19.18% of protein-centered variance, respectively. The exploratory global comparison yielded pseudo-*F* = 5.3106, *R*² = 0.6390, and allocation *p* = 6/1,680 = 0.00357. Root-mean-square (RMS) log2 centroid distances were 0.8644 for control/LPS, 0.5530 for LPS/treatment, and 0.9842 for control/treatment. Treatment was therefore closer to LPS than to control and remained farther from control than LPS was in this representation. The CLR global comparison also yielded *p* = 0.00357, with *R*² = 0.6604 and corresponding distances of 0.8426, 0.5305, and 0.9835. The global tests do not establish every pairwise difference and may reflect dispersion as well as centroid location. Neither distance nor variance explained measures injury or benefit.

Within-cohort sensitivity distinguished stable effect directions from retention of a significance threshold. Across the six relevant single-column omissions, all 13 candidates evaluated in the source omission analysis retained their treatment/LPS direction. However, PTGFRN retained *q* < 0.05 in none of the omissions and LAMB2 in two (Supplementary Table S3). Full-cohort Hedges *g* estimates were 11.19 for PTGFRN and 24.54 for LAMB2, reflecting in part small estimated pooled log2 SDs of 0.277 and 0.0858, respectively. These large estimates do not establish population reproducibility. The omission analyses reused overlapping samples and added no independent evidence.

### 3.6 Conditional calibration of aggregate reversal and profile geometry

The deposited LPS/control candidate set contained 227 nominal changes, of which 120 (52.9%) moved in the opposing direction after treatment and 119 (52.4%) became closer to control [2]. Conditional calibration repeated the initial screen in every allocation of the six LPS/treatment columns while holding the control columns fixed (Figure 5A; Table 4). Candidate counts ranged from 182 to 344, confirming that candidate selection varied under the reference procedure.

**Table 4.**
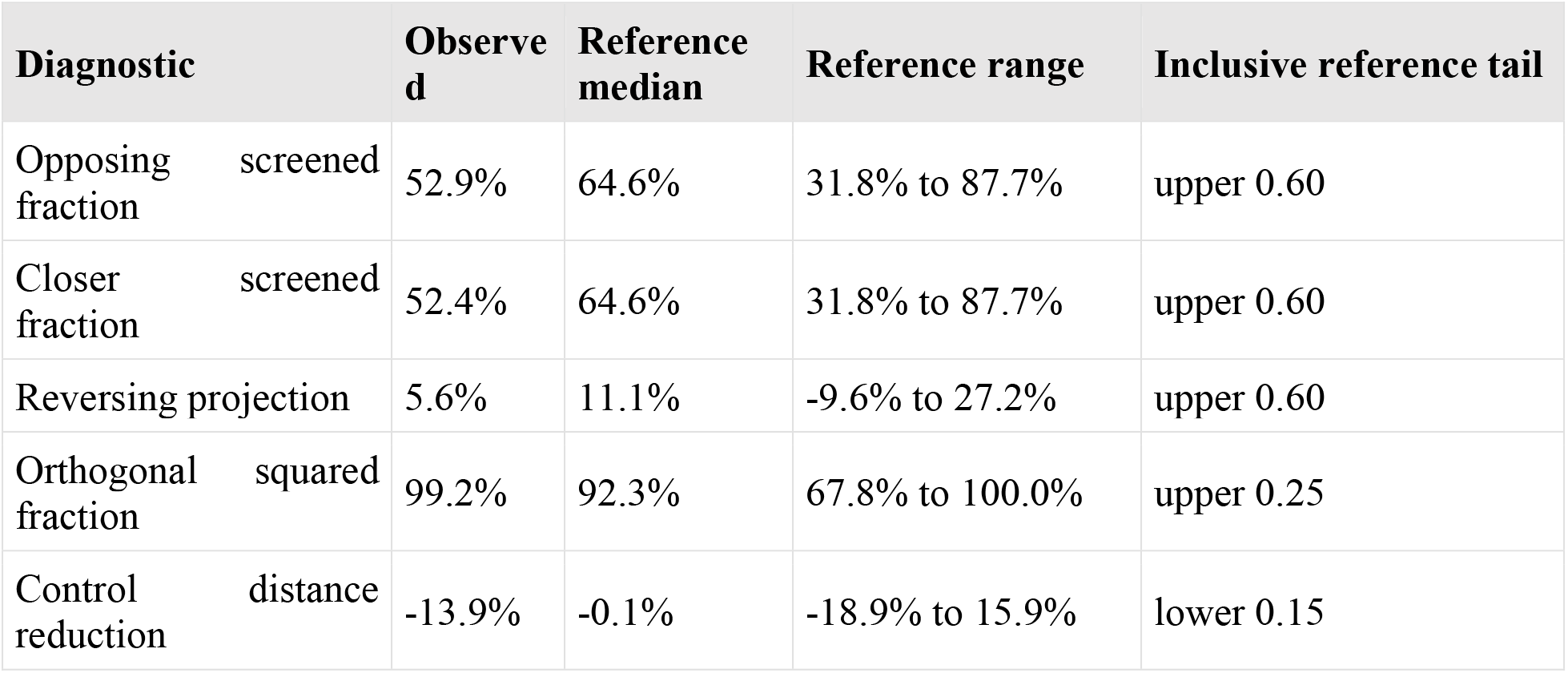
Complete selection conditional calibration All 20 allocations divide the six LPS/treatment columns into groups of three, with F7-F9 controls fixed. The initial screen is repeated under each allocation. Tails are inclusive count/20 reference fractions requiring exchangeability. These descriptive diagnostics do not estimate the biological fraction of recovery or constitute independent replication.

Across the 20 allocations, the median opposing fraction was 64.6% (range, 31.8-87.7%). Twelve allocations met or exceeded the observed 52.9%, yielding an inclusive upper reference fraction of 0.60. The closer-to-control criterion had the same upper reference fraction. Neither observed aggregate fraction was unusual under this conditional exchangeability reference. These results do not estimate how many individual changes arose from shared-reference sampling error or negate protein-specific formulation associations.

Conditional calibration also qualified the profile geometry. The observed reversing projection coefficient was 0.0564, and the component orthogonal to the LPS/control contrast represented 99.22% of squared treatment/LPS displacement. Under the conditional reference, the median orthogonal fraction was 92.25%, and five of 20 allocations reached or exceeded the observed fraction (inclusive upper reference fraction, 0.25). The primary control-distance reduction was - 13.86%, indicating a greater treatment/control distance than LPS/control distance. Exploratory within-group bootstrap percentile limits ranged from -30.74% to +6.92%. Three of 20 conditional allocations had an equally or more negative distance-reduction value (inclusive lower reference fraction, 0.15). These quantities describe the measured relative profile; they do not quantify recovered proteins or establish a distinct treatment-induced biological program (Figure 5B; Supplementary Figure S1C).

### 3.7 Morphology and proteomics characterized different aspects of the preparations

The combined comparison distinguished three descriptive patterns (Table 5). In kidney preparations, treatment circularity was closer to control while CQD remained elevated. Brain outline metrics changed little and were sensitive to field aggregation, while the brain proteome retained substantial group differences. Within that proteome, PTGFRN and LAMB2 had higher relative representation alongside persistent treatment/control differences in other proteins. These findings support interpretation at the level of each measured endpoint. They do not establish biological uncoupling of particle shape and protein composition, because the individual TEM-to-LFQ sample crosswalk was unavailable and the modalities sampled different components of the preparations.

**Table 5.**
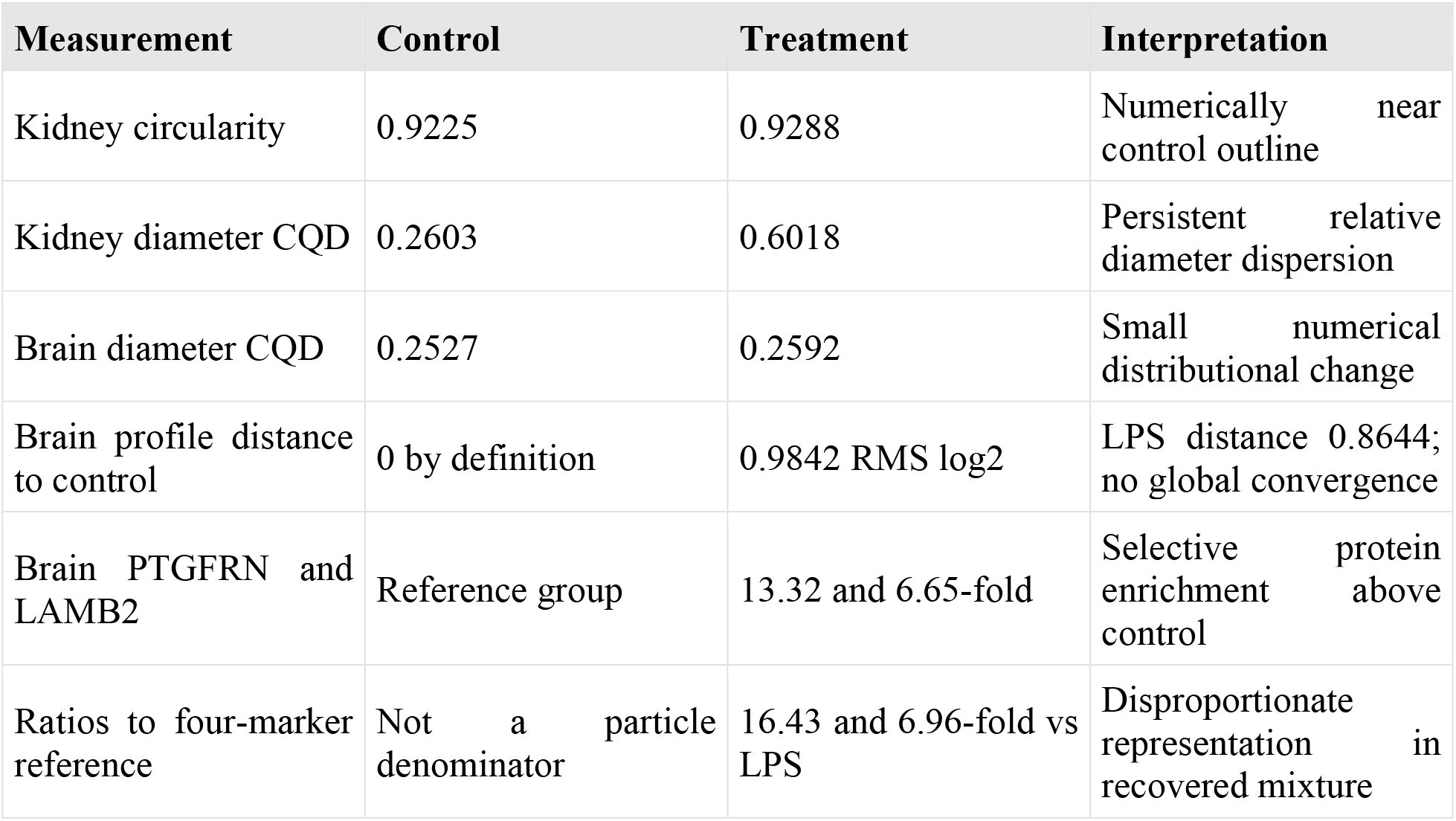
Comparison of morphology and protein composition TEM values are source-reported group means; proteomics values were recalculated. Endpoints differ in tissue coverage, denominator and sampling. They are compared descriptively and do not establish individual particle or animal correlations.

Missing specimen linkage constrained integration. In the source assignment analysis, all audited centered correlations admitted both signs across 216 possible within-group brain LFQ-TEM joins. For example, the correlation range was -0.782 to +0.618 for LAMB2/AR and -0.383 to +0.503 for PTGFRN/circularity (Supplementary Figure S2C). These ranges describe uncertainty in specimen identity, not biological associations. Treatment/LPS kidney-minus-brain point contrasts were +0.05737 for AR, +0.02200 for circularity, and +0.07934 for CQD. However, the envelope formed by the conditional interval endpoints across hypothetical common-animal pairings spanned zero for every endpoint. These source analyses could not be independently reproduced without the morphometry files. No kidney proteomics was available; the brain protein findings therefore cannot be assigned to renal EVs or used as a renal molecular explanation.

## 4 Discussion

This analysis identifies treatment-associated changes that differ across morphological and molecular endpoints. Kidney particle circularity was numerically close to control after treatment, while relative diameter dispersion remained elevated. Brain preparations showed selective protein enrichment without demonstrated convergence of the overall protein profile toward control. Identification recurrence provided a third perspective, describing consistency of detection across preparations. Together, these observations support a narrower interpretation than generalized recovery: treatment was associated with changes in the composition and measured properties of the recovered preparations, whose biological significance depends on particle identity, protein localization, and function. The proteomic cohort and leading protein discoveries were reported by our team [2]; the present contribution is their comparison with morphology and the additional assessment of measurement consistency and analytical sensitivity.

The kidney findings illustrate why average outline shape and distribution width should be interpreted separately. Treatment circularity differed from control by only +0.0063, whereas the coefficient of quartile dispersion (CQD) remained +0.3415 above control. The corresponding mean upper-to-lower quartile ratio (Q3/Q1) was 4.40 after treatment versus 1.79 in control. Thus, the measured fields could show similar average circularity despite broader relative diameter distributions. This pattern is compatible with changes in the proportions of EV subpopulations, membrane fragments, or aggregates, as well as differences among sampled grid regions. The available measurements cannot distinguish these explanations, but they show why a near-control mean circularity is insufficient to describe the preparation as morphologically restored.

Brain CQD varied over a narrower numerical range, and aspect ratio (AR) showed field-dependent trajectories. These observations require caution in cross-tissue interpretation because kidney samples contributed one measured field each, whereas brain samples contributed two to four. Unequal imaging coverage changes how well each sample summary represents grid heterogeneity. Moreover, hundreds of traced outlines remain nested within three biological preparations per group. Increasing the number of particles measured in the same selected fields does not increase biological replication. Because the primary morphological endpoints did not remain significant after correction for multiple comparisons, the results support descriptive patterns rather than established group differences; they also do not establish equivalence.

Negative-stain TEM records projected particle profiles after adsorption, staining, and drying. Cryo-TEM has documented substantial EV architectural diversity across sources and separation methods [16], and Rikkert and colleagues showed that operator-selected images can conceal preparation differences apparent at predefined grid locations [17]. Roundness therefore cannot serve as a universal measure of vesicle integrity or function. AR, circularity, and within-field CQD are unaffected by a uniform change in scale, permitting limited comparisons while absolute calibration remains unresolved. They remain sensitive to image resolution, segmentation, tracing, and field selection. Unreconciled scale records prevent reliable absolute-diameter claims, and the absence of original measurement rows and regions of interest from the records available for this reanalysis prevents independent reproduction of the particle-level results.

The molecular findings distinguish treatment-associated enrichment from movement toward control. PTGFRN and LAMB2 were already close to the control mean after LPS and increased markedly in the treatment group. Their trajectories therefore describe a treatment-associated composition rather than correction of a clear LPS-induced deficit. C3 and WFS1 followed different numerical patterns, moving toward control after treatment. Meanwhile, numerical treatment/control differences persisted for MX1, AQP4, SLC1A2, GRIA4, and MFGE8. Selected proteins approaching control consequently cannot establish recovery of the preparation as a whole. The supported interpretation is selective remodeling of relative protein representation in the recovered material.

The sensitivity analyses clarify the strength and scope of this interpretation. PTGFRN and LAMB2 remained the only primary treatment/LPS discoveries under a stricter unique-peptide eligibility criterion. This finding reduces concern that their discovery status depended solely on the original unique-plus-razor peptide filter. However, changing the eligible protein set also changes the Benjamini–Hochberg testing family; it does not independently verify peptide identity or reproduce the biological effect. Likewise, the post hoc marker-reference ratios showed that their enrichment exceeded the behavior of the measured CD9, ALIX, FLOT1, and FLOT2 panel. A common multiplicative sample factor cancels from these ratios, but protein-specific measurement effects, changes in the reference proteins, and differences in the recovered particle mixture remain possible. Loss of Benjamini–Hochberg significance after centered log-ratio transformation further indicates that statistical support depends on the analytical representation [8]. Large observed relative effects therefore warrant validation without implying precisely established population effects.

The aggregate recovery analysis addresses a separate statistical issue. The LPS mean appears in both the LPS/control and treatment/LPS contrasts, with opposite signs. Sampling error in that shared mean can therefore make the two contrasts appear oppositely directed, especially when proteins are first selected for an LPS/control difference. The complete-selection calibration repeated both protein selection and trajectory assessment in each allocation. The observed fraction of opposing trajectories did not exceed its conditional reference distribution, providing no support for an aggregate recovered-protein count under that procedure and its exchangeability assumption. This result does not exclude individual treatment responses or determine how much regression to the mean contributed. Similarly, a component perpendicular to the LPS-associated direction describes the geometry of the protein profile without identifying an alternative biological program.

These distinctions also explain why TEM and proteomics need not show parallel patterns. TEM characterizes selected projected outlines, while bulk proteomics measures proteins across the injected digest, including material that may not be represented by traced particles. Substantial compositional differences can therefore coexist with small outline changes. Their joint analysis reveals complementary properties of the preparations, but does not demonstrate molecular and morphological coupling within individual EVs.

The treatment group had the largest fraction of proteins detected in every preparation within the group and the highest average within-group overlap. This ordering persisted with a stricter per-run peptide threshold, supporting its reproducibility within the supplied identification exports. Its biological interpretation is less certain. In data-dependent acquisition (DDA), recurrence reflects protein abundance together with precursor selection, identification confidence, and protein inference. LPS identification depth varied by nearly 400 eligible accessions between preparations, illustrating how differences in analytical coverage can influence overlap.

The 267 accessions identified in all nine individual lists and the 953 protein groups with complete quantitative profiles in the combined matrix represent different evidence units. Separate identification searches, combined protein grouping, and quantitative signal extraction need not produce identical coverage. The available records do not establish which upstream operations account for the discrepancy. DDA sampling and quantification-centered workflows explain why a missing run-specific tandem mass spectrum does not necessarily indicate absence of the corresponding peptide ion [22,23]. Accordingly, identification indicators and peptide-spectrum match counts were not substituted for label-free quantification (LFQ) intensities when interpreting abundance changes.

A larger recurring core therefore indicates more consistent detection in these measurements, without establishing reduced EV heterogeneity, healthier tissue, or a more stable secretory program. Similarly, sample-accumulation curves describe how the observed protein union expands as preparations are combined. With three preparations per group, they cannot establish proteome completeness or separate biological diversity from analytical undersampling. Recurrence is most useful here as a complementary measure of coverage and a guide to targets requiring peptide-level confirmation.

PTGFRN offers a specific starting point for interpreting the selective enrichment. Its association with CD9 and CD81 in cellular systems [27], together with its use as a scaffold for high-density EV surface display [9], makes altered representation of PTGFRN-containing membrane assemblies a plausible hypothesis. Its enrichment relative to each measured reference protein strengthens the case for examining this candidate beyond a change in total preparation protein. Nevertheless, equal-protein-input LFQ and marker ratios cannot distinguish more PTGFRN per particle from a greater proportion of PTGFRN-positive particles, altered source-cell contributions, or increased recovery of other PTGFRN-containing membranes. They also do not establish targeting competence.

LAMB2 extends this question to extracellular-matrix association. Its disproportionate recovery relative to the same marker panel could reflect vesicle association, adsorption to a protein corona, matrix material released during tissue processing, or other co-separated extracellular components. Studies of tissue-specific rescue in Lamb2-null mice illustrate that laminin β2 function depends on its anatomical localization [10]. Consequently, enrichment of PTGFRN and LAMB2 in the same bulk preparation motivates a membrane–matrix association hypothesis, while leaving their physical relationship unresolved. They have not been shown to occupy the same particles, interact directly, or mediate repair. Because LAMB2 abundance was measured only in brain preparations, its established renal functions cannot be invoked to explain the kidney morphology findings without corresponding renal measurements.

The residual protein differences broaden the compositional question without resolving cellular origin. MX1 is consistent with an interferon-associated component, whereas AQP4 and SLC1A2 are associated with astroglial membrane physiology and GRIA4 with excitatory receptor biology. Their representation in an EV-enriched fraction may change through altered tissue expression, protein sorting, source-cell contributions, or membrane recovery. A lower fraction-associated AQP4 or SLC1A2 signal does not directly measure brain water homeostasis or glutamate transport. Matched source-tissue and fraction measurements are needed to distinguish tissue-level changes from redistribution into recovered extracellular material. The different tissue and brain-EV protein profiles reported after acute LPS exposure by Vassileff and colleagues provide context for this distinction, although their nominal-significance screening results and different experimental model do not independently validate the present candidates [4].

Administration of an EV-based formulation introduces further possible sources of compositional change. Altered endogenous secretion, persistence of administered material, changes in clearance, and co-recovery of tissue components could all contribute. In a systemic LPS mouse model, Pavlova and colleagues reported changes in EV clearance and protein-corona composition, with limited functional cargo delivery despite increased uptake [29]. That model differs from the present intracerebroventricular challenge, but demonstrates why accumulation of EV-associated signal and productive cargo delivery require separate measurements. Without carrier tracing, unloaded-carrier controls, and loading-process controls, the present protein changes cannot be assigned specifically to the administered miRNAs.

ER and calcium-associated candidates remain secondary hypotheses. Mechanistic studies have linked WFS1 to ATF6α regulation and TMX1 to ER calcium handling through targeted perturbation and compartment-specific functional assays [11,12]. Their abundance trajectories in the present fractions therefore justify examining ER-related processes, but cannot establish reduced tissue ER stress, restored ER–mitochondrial calcium transfer, or improved respiration. Similarly, the known lipid-biochemical roles of CDS2 and AGPAT3 [13,14] provide a rationale for lipidomics, while their treatment/LPS q values of 0.349 and 0.398, respectively, do not establish treatment-associated restoration of membrane lipid composition.

C3 requires the same distinction between a descriptive trajectory and a demonstrated functional response. Its abundance changed from 2.053-fold relative to control after LPS to 1.068-fold after treatment, but the treatment/LPS q value was 0.412. Total C3 abundance cannot distinguish complement activation fragments, measure complement activity, or identify the C3-positive cellular state described in reactive-astrocyte studies [15]. These candidates are therefore most useful for defining focused validation questions, rather than expanding the interpretation into multiple established pathway effects.

## 5. Limitations

This exploratory study is limited by the small male-only cohort, unequal TEM sampling, incomplete imaging records and calibration, unresolved sample linkage, and possible co-recovery of nonvesicular material. These constraints limit cross-tissue comparisons and individual morphology–proteome associations. Without carrier tracing, renal proteomics, and functional delivery measurements, the findings cannot establish brain-to-kidney EV transport or miRNA-specific therapeutic effects.

## 6. Conclusion and perspectives

This study integrates tissue particle morphology with reanalysis of brain proteomics to characterize EV-enriched preparations following LPS challenge and miRNA-associated vesicle administration. The principal observation is that treatment-associated patterns differed across measured endpoints. Kidney circularity was numerically close to control while relative diameter dispersion remained elevated. Brain preparations showed selective protein enrichment without demonstrated convergence of the overall protein profile toward control. Together, these findings emphasize that average particle shape, size dispersion, and protein composition capture different properties of recovered preparations and should be interpreted together when evaluating treatment responses.

The reanalysis reinforced PTGFRN and LAMB2 as priorities for further investigation. Both retained their primary treatment/LPS discovery status under stricter unique-peptide filtering, and post hoc marker-reference ratios supported their disproportionate representation relative to the measured marker panel. These observations motivate a focused hypothesis concerning changes in membrane- and matrix-associated material. Determining whether these proteins identify particular vesicle populations, associate with vesicle surfaces, or accompany co-separated components would clarify the biological meaning of their enrichment. The contribution extends the previously reported proteomic cohort by relating its findings to morphology and additional assessments of detection consistency and analytical sensitivity.

For exosome research, the broader implication is the need to connect structural characterization with molecular localization and biological activity. A preparation with a control-like average outline may still differ in particle distribution or protein composition. Likewise, enrichment of a membrane-associated protein does not specify its abundance per vesicle or its contribution to cargo delivery. Evaluating these properties separately provides a clearer basis for identifying which preparation characteristics are relevant to biological responses.

Future studies should prioritize an independent cohort with verified animal identities, calibrated imaging, and systematic field sampling. Targeted peptide measurements in matched tissue, administered formulations, and EV-containing fractions should be combined with immunocapture, single-particle localization, and topology-appropriate protease assays. Reporting analytes per tissue input, preparation protein, and particle number would help distinguish total recovery from relative representation. Prospectively defined protein ratios or functional-module balances could provide focused molecular endpoints, provided their membership, peptide coverage, and reference stability are established [24,25].

The next biological step is to connect these compositional findings with productive cargo delivery and relevant outcomes. For miRNA-associated interventions, this requires verified loading and RNA protection, delivery to the appropriate recipient compartment, and target engagement, supported by unloaded-vesicle and loading-process controls. Matched brain and kidney measurements over a justified time course could clarify tissue-specific responses, while inflammatory, neuronal, and renal assays would determine their functional relevance. This progression would help identify which structural and molecular attributes support reproducible evaluation of exosome-based interventions.

## Supplementary Methods

### S1. Effect estimation distribution shape and specimen assignment

For an accession, Δ is the difference in mean log2 abundance, 2^Δ^ the geometric-mean fold change and V_A_ and V_B_ the unbiased group variances. At n_A_ = n_B_ = 3, Welch standard error and degrees of freedom are [20]:

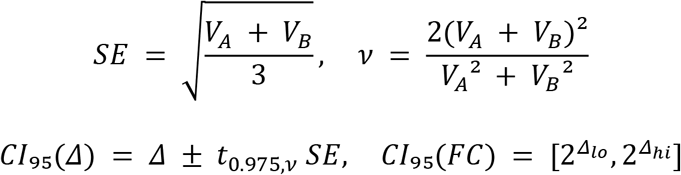

Pointwise 95% intervals used fractional Welch degrees of freedom without integer rounding. Such intervals were already part of the deposited analysis [2]; the present audit specifies their calculation and uses the exact gamma correction for Hedges g [18]. They are normal-model intervals, not simultaneous coverage statements or substitutes for BH adjustment. The pooled log2 SD is sp = √((VA + VB)/2), with pooled degrees of freedom νp = 4:

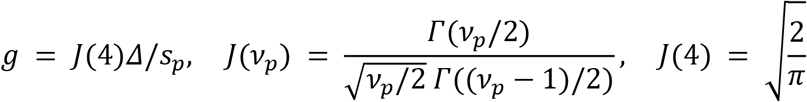

The usual bias-correction interpretation assumes normal populations with common variance; under unequal population variances, g remains a descriptive pooled-SD standardization. No g significance threshold or confidence interval was inferred. A fixed-feature centered-log-ratio (CLR) sensitivity changed the reference to each sample’s geometric mean of the same 953 measured proteins [19].

The source-reported primary treatment/LPS pointwise intervals were 14.71-fold (9.49-22.78), g = 11.19 for PTGFRN and 6.23-fold (5.42-7.16), g = 24.54 for LAMB2. Their pooled log2 SDs were 0.277 and 0.0858. These extreme standardized estimates reflect small estimated cohort variances. C3 and WFS1 pointwise intervals were 0.520 (0.356-0.760) and 2.79 (2.01-3.87), although neither met the primary 953-test BH threshold.

Within each field, Q1, Q2 and Q3 were linearly interpolated ECD quartiles on the original recorded diameter scale, then transformed to Lk = log2(Qk). We applied the Bowley/Galton quartile form to those transformed quartiles [21]. This compares upper and lower multiplicative half-interquartile ranges and adds central-distribution shape beyond CQD.

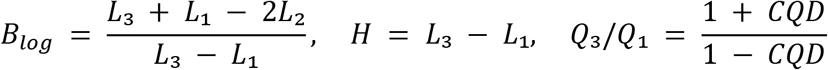

In the source morphology analysis, a preparation’s asymmetry was the median of its field coefficients. Positive Blog means Q3/Q2 exceeds Q2/Q1; negative values mean the reverse. H and Q3/Q1 are monotone re-expressions of field CQD, without new hypothesis tests. Nonlinear transforms and aggregation over even field counts or sample means do not generally commute. The two new exploratory asymmetry omnibus tests used all 1,680 sample allocations with Holm correction across two tissues; four planned contrasts used 20 allocations and Holm across four tests. These are additions developed during secondary analysis, not prospectively specified confirmatory outcomes. One-outline omissions assessed sampling sensitivity, with ranges reported separately from sampling intervals.

For dimensionless endpoint y and groups g,h, a tissue contrast-of-contrasts directly compared kidney (K) and brain (B) response magnitudes. Its point estimate does not depend on within-group animal correspondence.

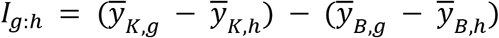

In the source pairing sensitivity, conditional on the same animals supplying both tissues, all (3!)³ = 216 within-group pairings were enumerated; each two-group contrast has only 36 distinct relevant pairings. For each join π, three paired tissue differences di,g = yK,π(i),g − yB,i,g gave variance Vd,g,π = VK,g + VB,g − 2CovKB,g,π and SEI,π = √((Vd,g,π + Vd,h,π)/3). Fractional-df Welch intervals used these paired-difference variances [20]. Minimum/maximum SE and the union of interval endpoints describe dependence on the hypothetical mapping. The union is not itself a calibrated 95% interval; no pairing, smallest interval or independence model was adopted as biological truth. Different field coverage remains inseparable from tissue-response interpretation.

In the source identifiability audit, the analysis enumerated all 216 possible within-group brain LFQ-TEM joins for PTGFRN, LAMB2, C3 and WFS1 against three equal-field morphology endpoints. Within-group centered Pearson correlations were recalculated for each join. The minimum-maximum envelope describes dependence on unresolved specimen identity, not a confidence interval or probability over joins. No positional join was treated as observed truth, and no biological association test was inferred from an arbitrary pairing. With centered variables, averaging over all within-group permutations forces the mean covariance to zero; both-sign envelopes therefore chiefly demonstrate missing linkage, not evidence against an underlying association.

Source-reported conditional treatment/LPS kidney-minus-brain assignment ranges were −0.1211 to +0.2358 for AR, −0.02659 to +0.07059 for circularity and −0.3146 to +0.4732 for CQD. They are unions of interval endpoints across hypothetical joins and have no calibrated 95% coverage. Within a kidney control field, a log-quartile asymmetry estimate of 0.9044 changed to 0.3095-0.6233 after individual profile omissions, illustrating sparse-field sensitivity.

### S2. Framework for future functional module validation

This section specifies future functional-module validation. The completed descriptive marker-reference ratios in Section 2.5 use the same log-ratio principle without functional-module inference. Let A and B be fixed, externally defined, disjoint modules of sizes a and b. For positive measured intensity I in each animal s, a module balance is [24]

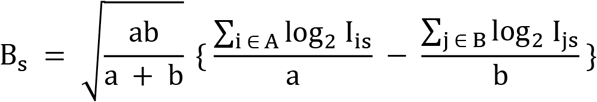

One animal contributes one balance score. Membership must remain fixed; averaging only whichever proteins are detected in each sample changes the endpoint. A prespecified protein-pair contrast is

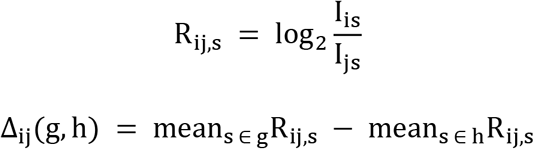

If Iis = cs ri Qis, the sample-wide multiplicative factor cs cancels within a balance. Stable protein response factors ri cancel from between-group differences, although raw cross-protein intensities are not molar proportions. These properties do not correct sample-dependent digestion, interference or changing protein inference. Three-versus-three two-sided exact label tests remain limited to 20 allocations and minimum P = 0.10. Independent validation, a small endpoint family and compartment-specific measurements are required before a functional interpretation [25].

---

**Figure S1.**
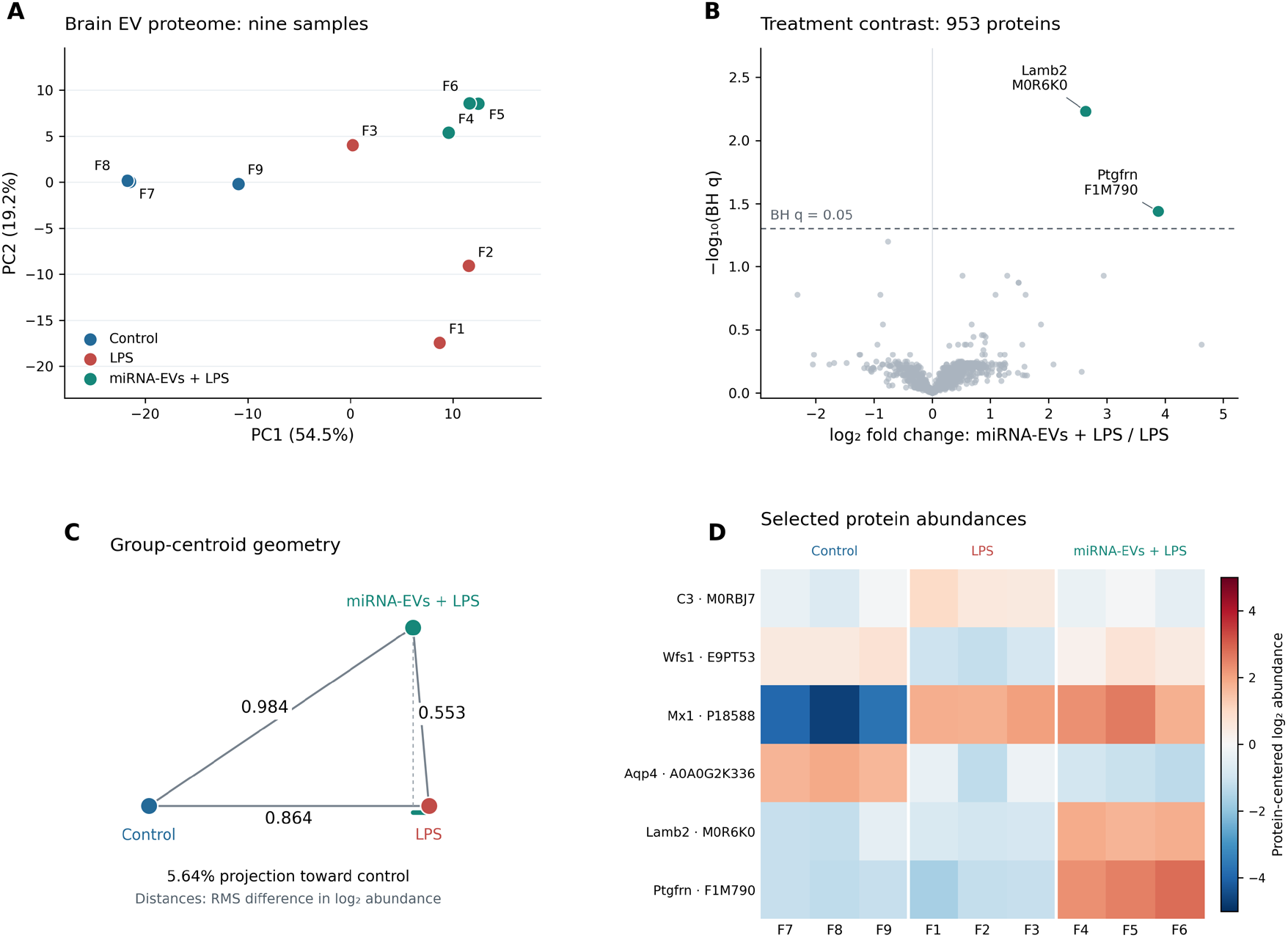
Reused brain proteomics reference results. (A) Protein-centered PCA of log2 abundance for the reused 953-feature matrix. F7-F9 are control, F1-F3 LPS and F4-F6 treatment; source-column identifiers do not establish individual TEM correspondence. (B) Treatment/LPS log2 fold change and contrast-specific BH q from Welch tests, n = 3 per group, adjusted across 953 proteins. PTGFRN and LAMB2 are the two primary discoveries. (C) Triangle preserving full-feature RMS log2 centroid distances. The reversing projection coefficient 0.0564 is not a fraction of recovered proteins. (D) Six selected accession-specific protein trajectories, centered across the nine samples; colors represent centered log2 values, not z scores. All panels summarize the same cohort and substantive proteomic results reported in Brown et al. [2], and are retained as context rather than new or independently validated findings. The current composite is not evidence of replication.

**Figure S2.**
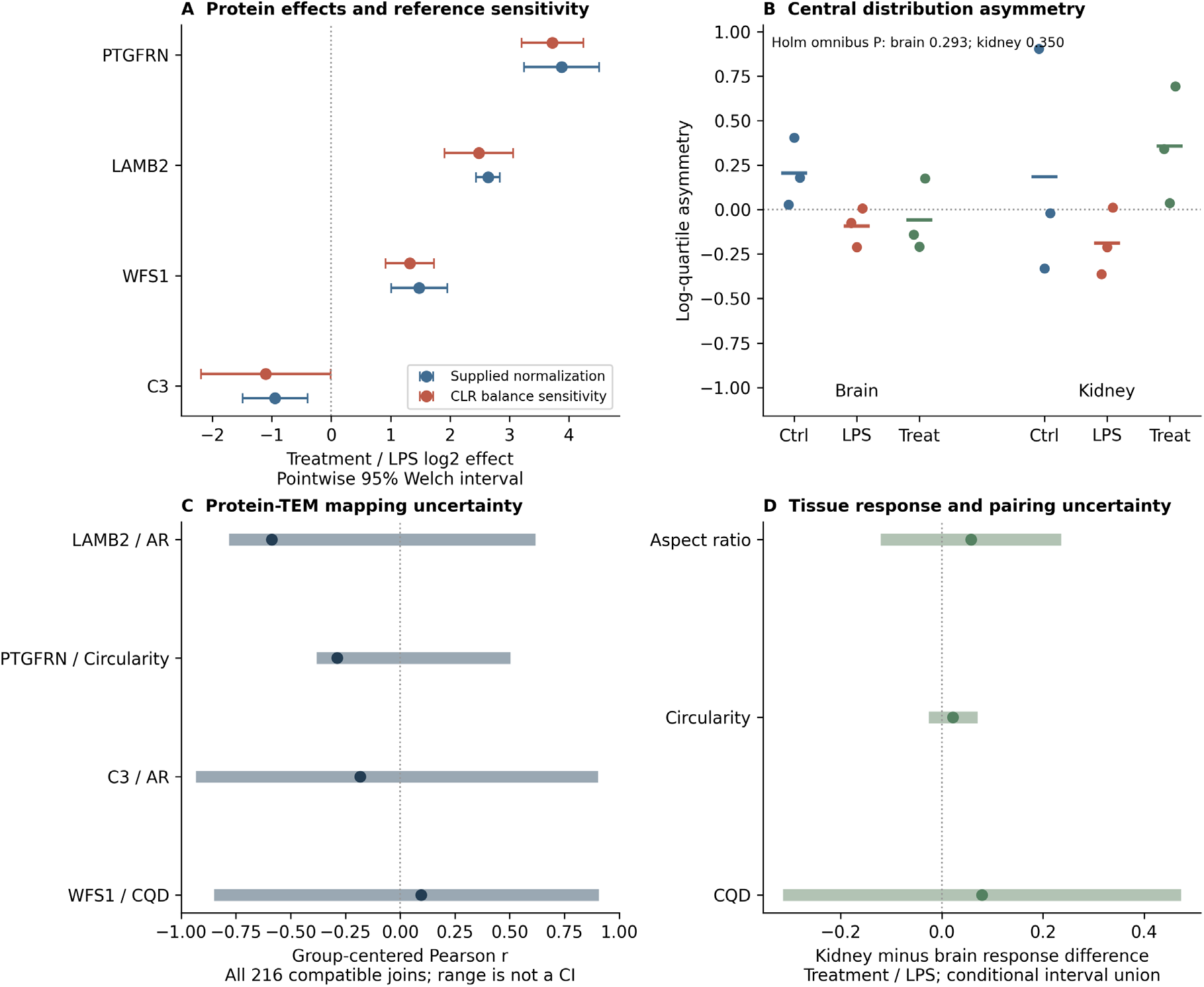
Effect estimation distribution shape and specimen assignment uncertainty. (A) Illustrative accession-specific treatment/LPS log2 effects under primary and fixed-953-feature CLR references with pointwise 95% fractional-df Welch intervals. These intervals are not adjusted for multiplicity; C3 and WFS1 are not primary BH discoveries, and neither PTGFRN nor LAMB2 retains treatment/LPS q < 0.05 under CLR. (B) Preparation-level median field log-quartile asymmetry, n = 3 per group, with group means indicated. Holm adjustment covers the two added exploratory tissue omnibus tests. (C) Full ranges of centered protein-morphology correlations across 216 possible within-group brain joins for four illustrative associations. Dark points show the unverified positional assignment solely for orientation; they are not verified sample correlations. The ranges are assignment envelopes, not confidence intervals or probabilities, and cannot test biological coupling. (D) Treatment/LPS kidney-minus-brain group contrasts. Bands span the union of conditional model-based 95% interval endpoints across hypothetical common-animal pairings, not a calibrated 95% confidence interval. Pairing is unverified and no favorable join is selected. These analyses add no biological replication.

**Figure S3.**
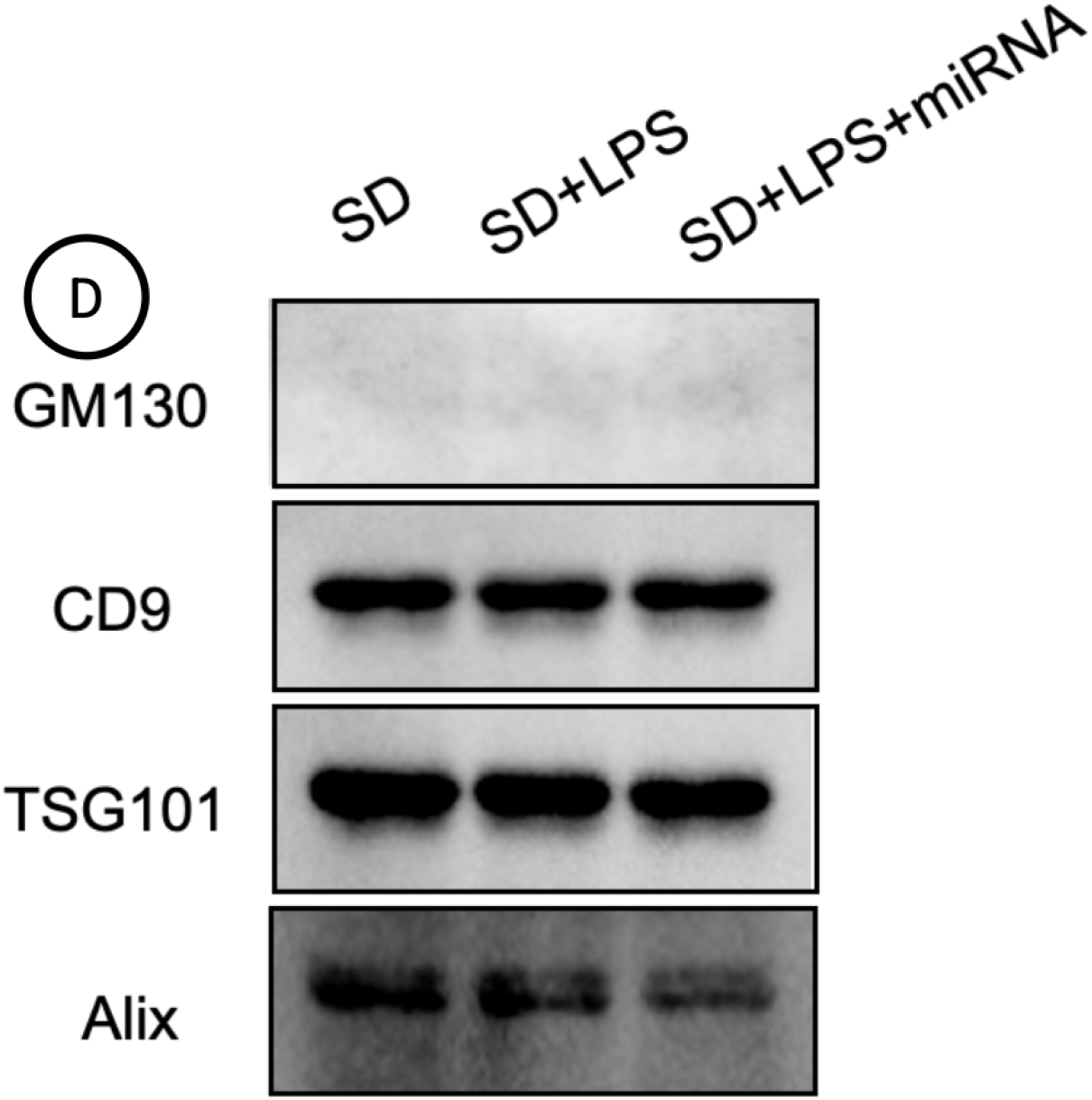
**Source supplied marker panel**. Source-supplied immunoblot panel displaying GM130, CD9, TSG101 and ALIX targets with lanes SD, SD+LPS and SD+LPS+miRNA. D is the original source panel designation. This panel was duplicated in both original tissue composites and is retained once without organ attribution. Larger source TIFF captures exist for several targets, but tissue identity, sample correspondence, loading, molecular-weight records and complete annotated membranes require confirmation. Signals are descriptive; the panel does not establish kidney-specific purity, quantitative treatment effects or absence of cellular material. Collagenase-associated proteolysis further limits interpretation of a weak GM130 signal [26].

**Figure S4.**
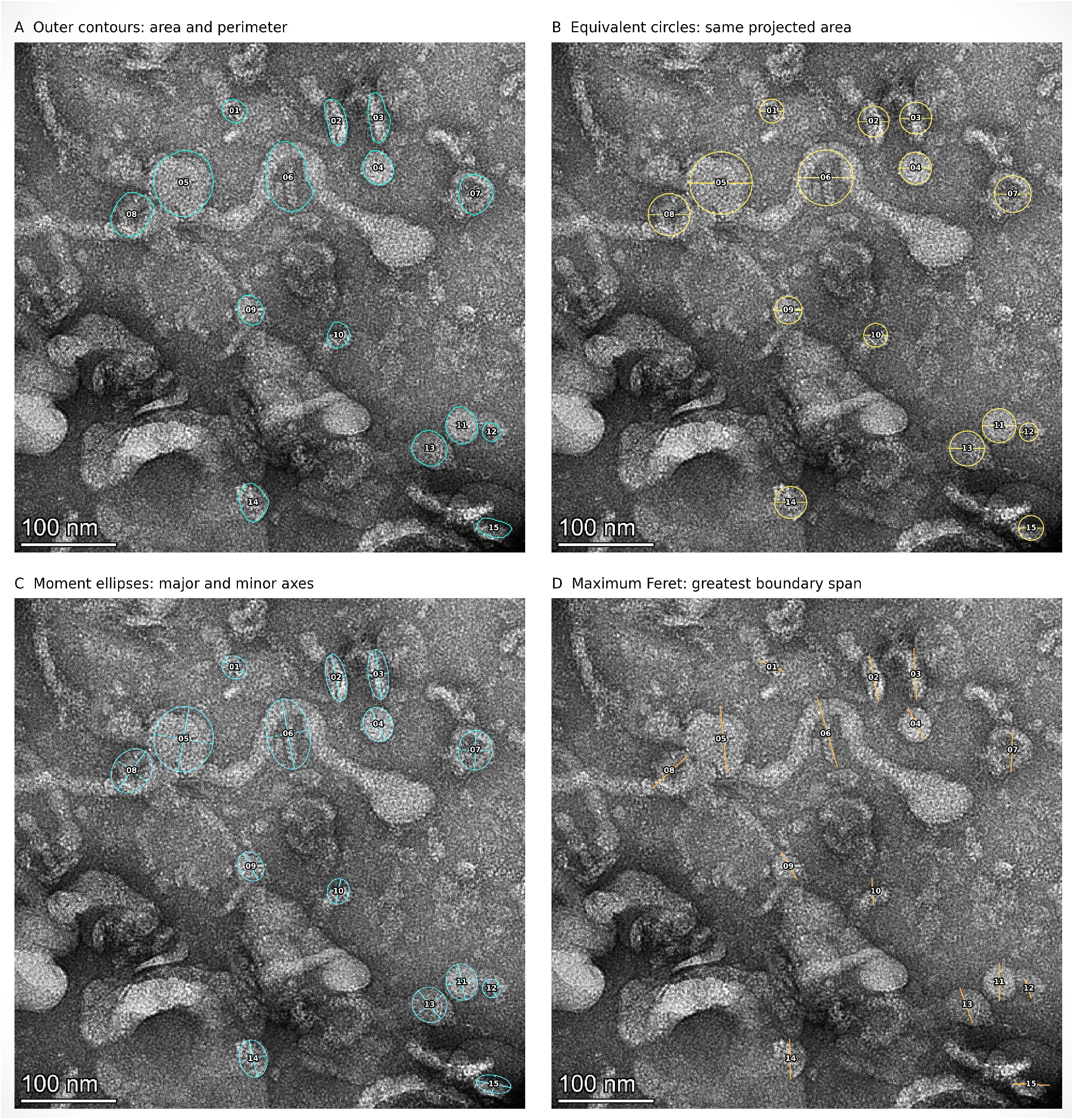
Illustration of contour derived measurements. Retained source illustration showing traced particle contours and derived measurements. These panels illustrate segmentation and measurement definitions; they do not add biological samples or independently validate the reported group summaries. Scale and annotation limitations discussed in Methods apply to physical-size interpretation.

**Table S1.**
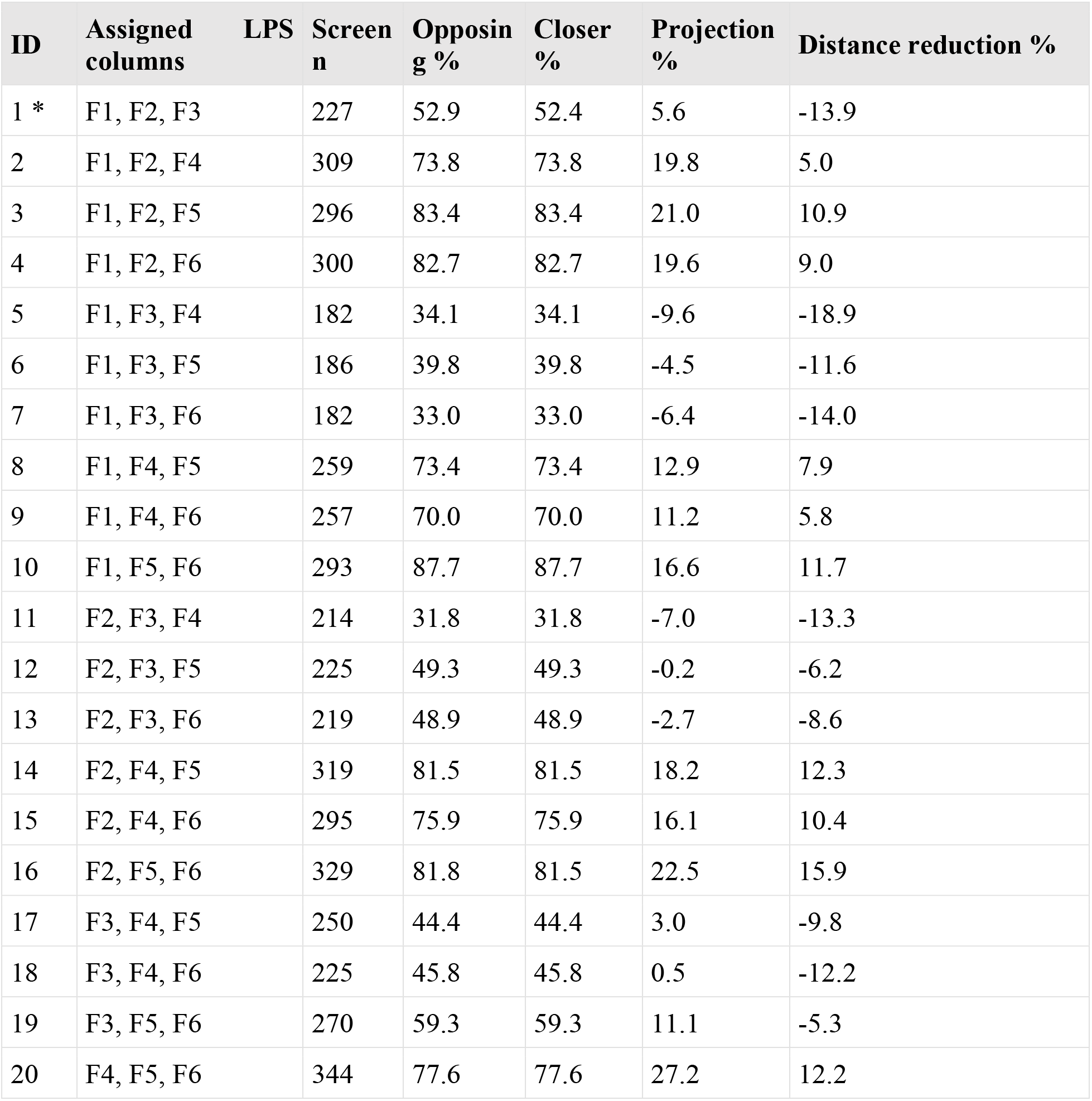
All conditional allocations. F7-F9 control columns are fixed. The assigned treatment group is the complement of the listed LPS group within F1-F6. Asterisk identifies the observed allocation. The initial candidate screen is repeated for each allocation; its size is not fixed at 227. All displayed values are descriptive reference results, not additional independent observations.

**Table S2.**
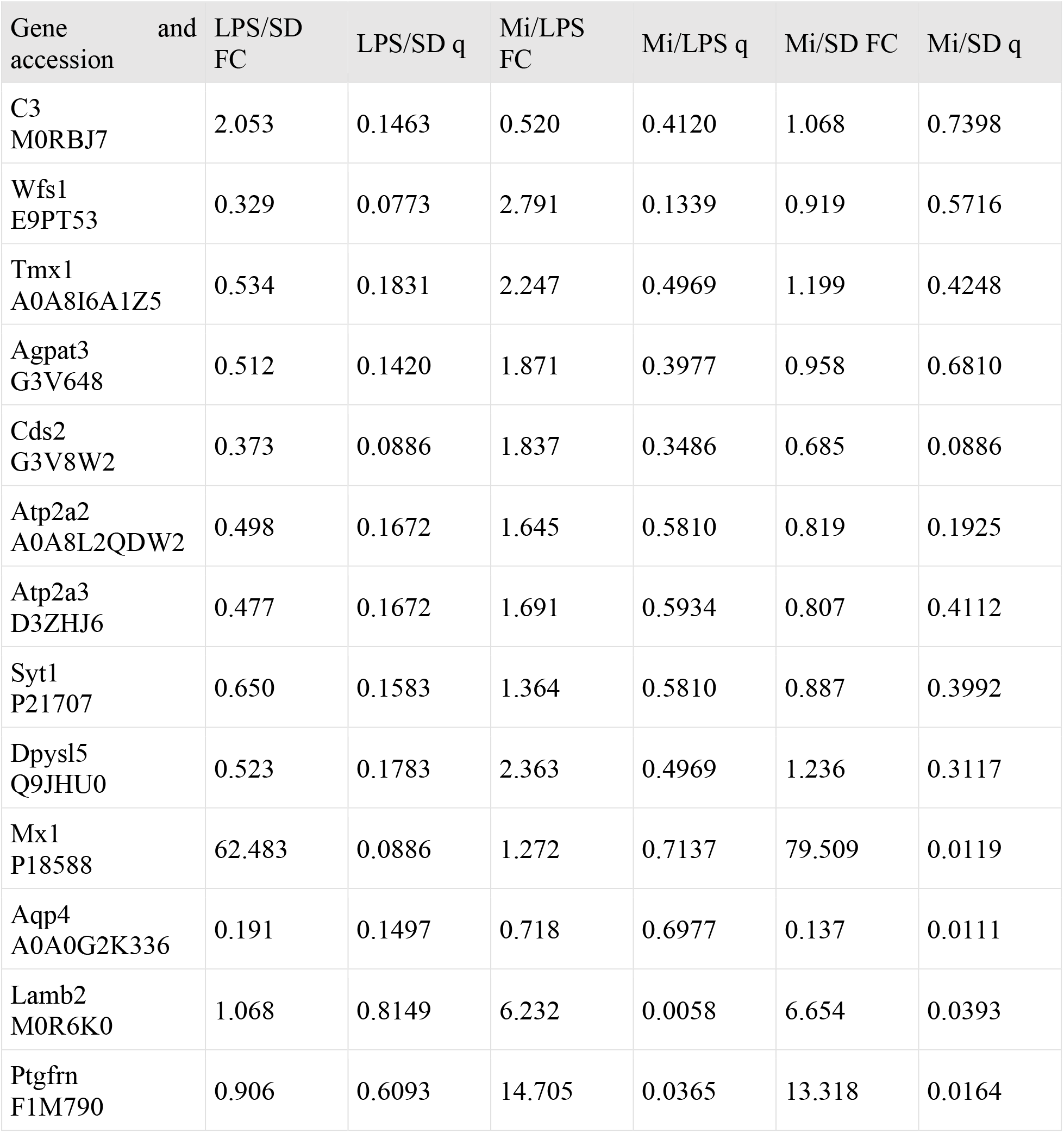
Reused selected primary protein trajectories Values recalculated from the supplied LFQ workbook for the deposited cohort [2], retained to interpret the secondary analyses. FC is the ratio of geometric means; q is BH-adjusted across 953 protein groups within its contrast. Exact accession identifiers are provided beside gene labels. Near-control means do not establish equivalence, and post hoc candidates are not independent validation.

**Table S3.**
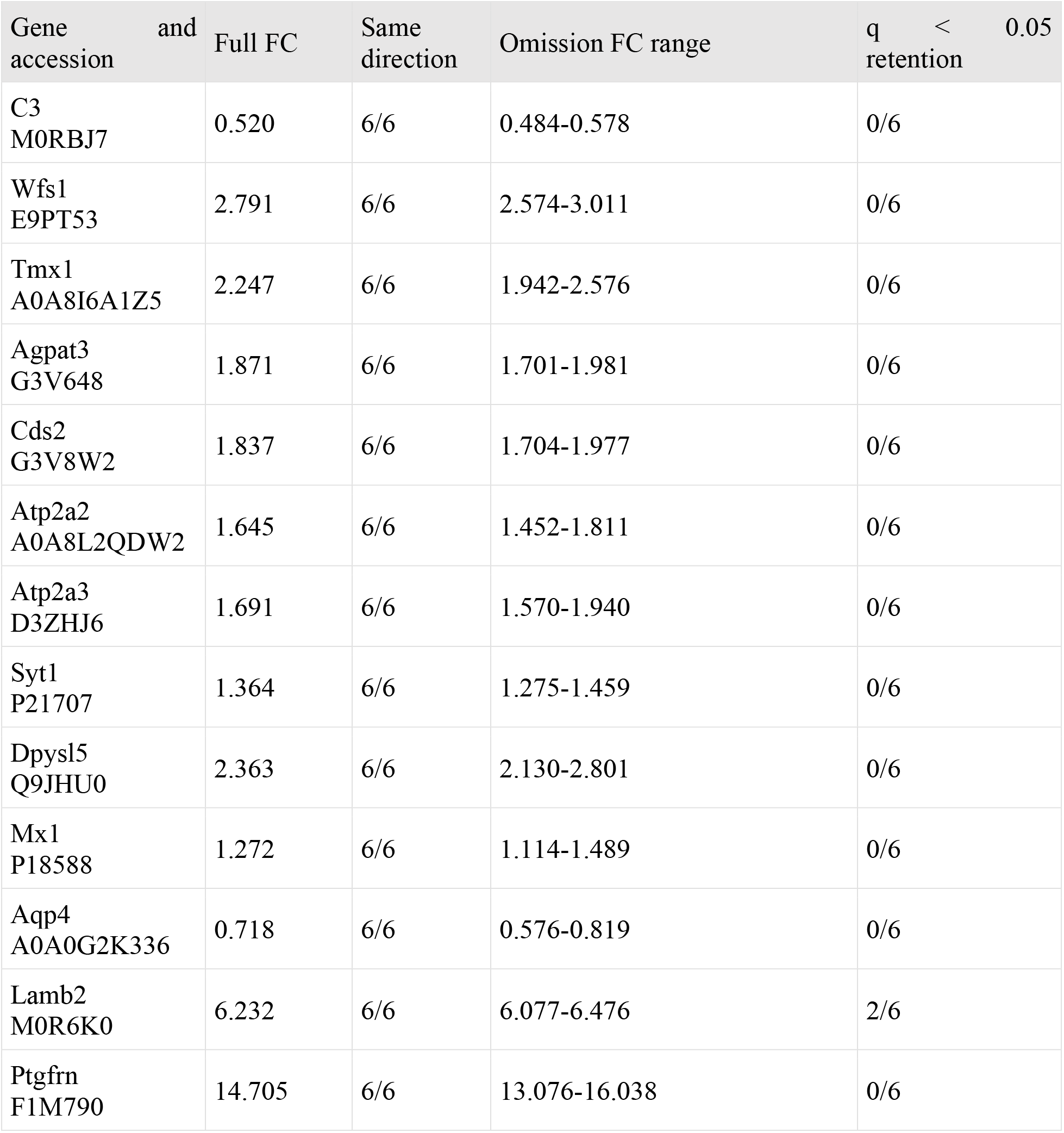
Expanded within cohort omission summary Expanded 13-protein summary recalculated from the supplied workbook using the omission procedure applied to the deposited cohort [2]. Six omissions affect treatment/LPS by reducing either comparison group to two samples; control omissions leave that contrast unchanged. Directional agreement is within-cohort stability, not independent replication; q-threshold retention at reduced n does not measure biological reproducibility.

**Table S4.** Exploratory marker reference ratios. Treatment/LPS ratios calculated within each sample before averaging on the log2 scale. All quantities are descriptive and post hoc. A common multiplicative sample factor cancels; these values do not measure molar stoichiometry or copies per EV. The last row uses the geometric mean of the four reference proteins. No new significance tests were applied.

| Reference | Reference FC | PTGFRN/reference FC | LAMB2/reference FC |
| --- | --- | --- | --- |
| CD9 A0A8I5ZZU3 | 0.865 | 17.01 | 7.21 |
| ALIX Q9QZA2 | 0.937 | 15.70 | 6.65 |
| FLOT1 Q9Z1E1 | 0.797 | 18.44 | 7.82 |
| FLOT2<br>A0A0G2K0Z1 | 0.993 | 14.80 | 6.27 |
| Four-marker<br>geometric mean | 0.895 | 16.43 | 6.96 |

**Table S5.**
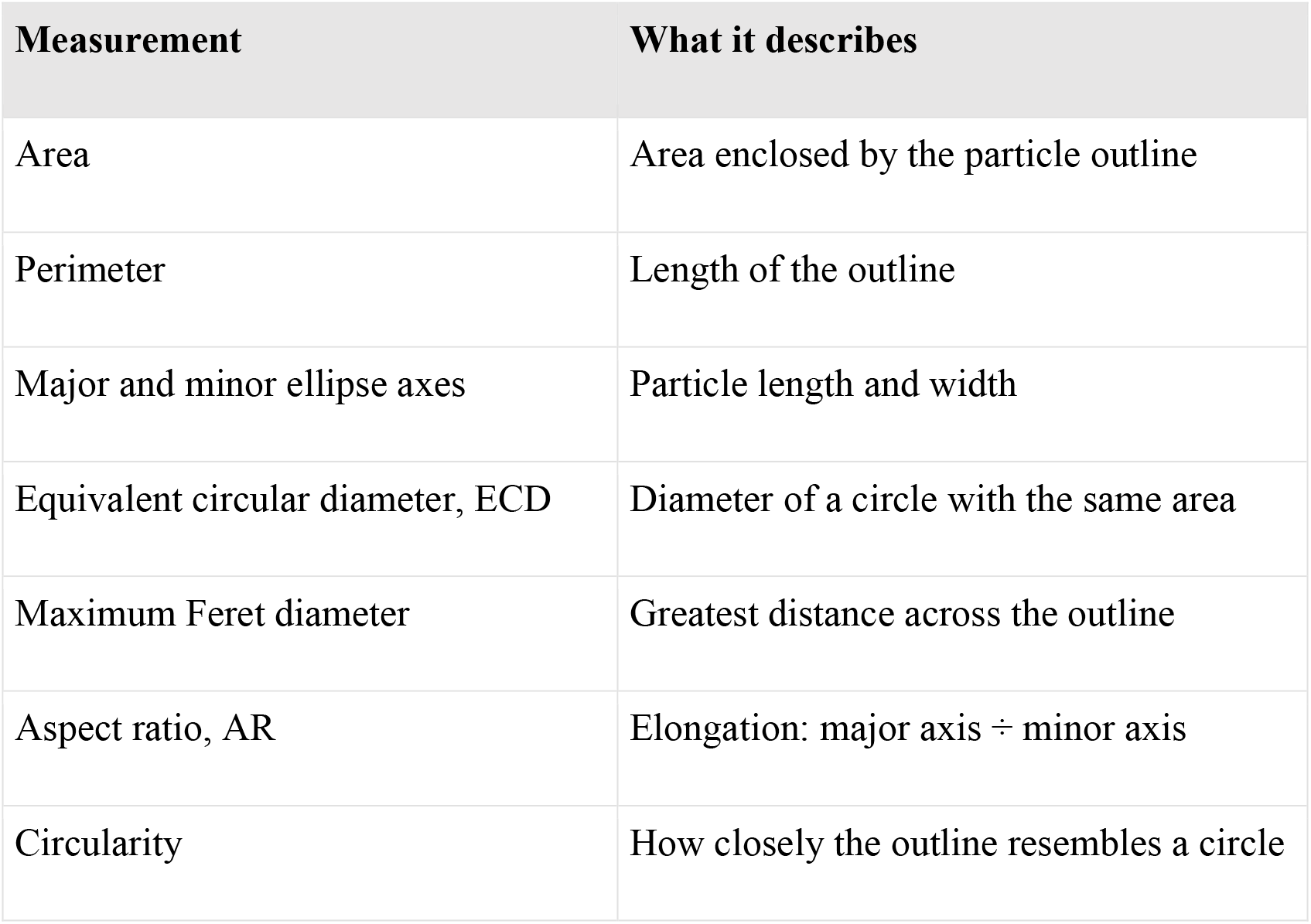
Definitions of contour measurements

| Measurement | What it describes |
| --- | --- |
| Area | Area enclosed by the particle outline |
| Perimeter | Length of the outline |
| Major and minor ellipse axes | Particle length and width |
| Equivalent circular diameter, ECD | Diameter of a circle with the same area |
| Maximum Feret diameter | Greatest distance across the outline |
| Aspect ratio, AR | Elongation: major axis ÷ minor axis |
| Circularity | How closely the outline resembles a circle |

